# Computational Modeling of Stapled Coiled-Coil Inhibitors Against Bcr-Abl: Toward a Treatment Strategy for CML

**DOI:** 10.1101/2023.11.15.566894

**Authors:** Maria Carolina P. Lima, Braxten D. Hornsby, Carol S. Lim, Thomas E. Cheatham

## Abstract

The chimeric oncoprotein Bcr-Abl is the causative agent of virtually all chronic myeloid leukemias (CML) and a subset of acute lymphoblastic leukemias (ALL). As a result of the so-called Philadelphia Chromosome translocation t(9;22), Bcr-Abl manifests as a constitutively active tyrosine kinase which promotes leukemogenesis by activation of cell cycle signaling pathways. Constitutive and oncogenic activation is mediated by an N-terminal coiled-coil oligomerization domain in Bcr (Bcr-CC), presenting a therapeutic target for inhibition of Bcr-Abl activity toward the treatment of Bcr-Abl+ leukemias. Previously, we demonstrated that a rationally designed Bcr-CC mutant, CCmut3, exerts a dominant negative effect upon Bcr-Abl activity by preferential oligomerization with Bcr-CC. Moreover, we have shown conjugation to a leukemia-specific cell-penetrating peptide (CPP-CCmut3) improves intracellular delivery and activity. However, our full-length CPP-CCmut3 construct (81 aa) is encumbered by an intrinsically high degree of conformational variability and susceptibility to proteolytic degradation, relative to traditional small molecule therapeutics. Here, we iterate a new generation of our inhibitor against Bcr-CC mediated Bcr-Abl assembly that is designed to address these constraints through incorporation of all-hydrocarbon staples spanning i, i + 7 positions in helix α2 (CPP-CCmut3-st). We utilize computational modeling and biomolecular simulation to design and characterize single and double staple candidates in silico, evaluating binding energetics and building upon our seminal work modeling single hydrocarbon staples when applied to a truncated Bcr-CC sequence. This strategy enables us to efficiently build, characterize, and screen lead single/double stapled CPP-CCmut3-st candidates for experimental studies and validation in vitro and in vivo. In addition to full-length CPP-CCmut, we model a truncated system characterized by deletion of helix α1 and the flexible-loop linker, which are known to impart high conformational variability. To study the impact of the N-terminal cyclic CPP toward model stability and inhibitor activity, we also model the full-length and truncated systems without CPP, with cyclized CPP, and with linear CPP, for a total of six systems which comprise our library. From this library, we present lead stapled peptide candidates to be synthesized and evaluated experimentally as our next-generation inhibitors against Bcr-Abl.

## Introduction

Protein oligomerization is pivotal for a diverse range of cellular processes, including signal transduction, gene expression, enzyme activity and regulation, and protein binding^1–15^. Such processes reflect but a fraction of the complete human interactome, comprised of an estimated 650,000 unique protein-protein interactions (PPIs)^16^ responsible for regulating physiological functions. Consequently, dysregulation within the interactome is also responsible for the development of disease states^17–20^. Indeed, aberrant oligomerization has been implicated in various pathological conditions and conformational diseases, such as diabetes mellitus^21, 22^, Alzheimer’s disease^23–25^, and Bcr-Abl^+^ cancers including chronic myeloid leukemia (CML) and acute lymphoblastic leukemia (ALL)^26–28^.

Approximately 95% of CML and 20% of ALL cases are characterized by the so-called Philadelphia Chromosome (Ph), a genetic abnormality which manifests due to a balanced reciprocal translation t(9;22) (q34;g11)^26–28^. In this event, the ABL1 gene of chromosome 9 is fused to the breakpoint cluster region (BCR) gene of chromosome 22, producing the BCR-ABL1 oncogene which encodes for the chimeric oncoprotein Bcr-Abl, a non-receptor tyrosine kinase with constitutive activity. Analogous to receptor tyrosine kinases, Bcr-Abl is activated by trans-autophosphorylation of tyrosine residues between kinase domains, clustered together upon homodimerization^29–31^. These phosphotyrosine residues then act as docking sites for SH2 domain substrates, activating RAS, PI3K/AKT, JAK/STAT, and MAPK signaling pathways which promote cell proliferation, inhibit apoptosis, and enhance cell motility—conferring growth advantage for leukemic cells^32–34^.

Bcr-Abl assembles as a dimer-of-dimers, mediated by an N-terminal oligomerization, or coiled-coil, domain of Bcr (Bcr-CC). Monomers associate as a dimer through formation of an antiparallel coiled coil and domain swapping, further associating as a tetramer through dimer stacking^29^. Bcr-CC (72 aa) comprises a short N-terminal helix (α1; residues 5-15) and a long C-terminal helix (α2; residues 28-67) connected by a flexible loop linker region. Helix α2 predominantly composes the dimeric interface, while helix α1 domain-swaps against helix α2 of the opposing monomer— packing the interface^29^. Deletion of Bcr-CC demonstrably diminishes transformation potential and interleukin-3 dependence imparted upon lymphoid cells^30^, while heterodimerization between Bcr-Abl and an isolated coiled-coil domain diminishes transformation and abrogates transphosphorylation^35–37^. Collectively, these findings substantiate Bcr-CC as an integral feature for leukemogenesis and highlight the dimeric interface as a therapeutic target for treating Bcr-Abl^+^ leukemias.

Previously, we designed and developed a dominant-negative Bcr-CC mutant (CCmut3) through a combinatorial approach utilizing biomolecular simulation, rational mutagenesis, and *in vitro* experimentation^38,39^. CCmut3 incorporates a set of six helix α2 mutations (K29E, C38A, S41R, L45D, E48R, Q60E) (**Figure 1**) which promote formation of preferential heterodimers with wild-type Bcr-CC while dissuading formation of deleterious CCmut3 homodimers (**Figure 2**). This CCmut3 construct demonstrably reduces transformation and proliferation, induces apoptosis, and diminishes phosphorylation of Bcr-Abl and its substrates in Bcr-Abl^+^ cells, including those with the clinically significant T315I point-contact mutant and the T315I/E255V compound mutant^38–40^. In a successive design iteration, a leukemia-specific cell penetrating peptide (CPP) was conjugated to the N-terminus of CCmut3 to improve intracellular delivery and activity (CPP-CCmut3)^40, 41^. This CPP (CAYHRLRRC), characterized by an arg-rich cell-penetrating motif (RLRR) and a lymph node-homing motif (CAY), was discovered by phage display, conferring greatly improved binding of directed phage to a panel of leukemia/lymphoma cells and patient samples compared to non-leukemic cells^41^. Indeed, we have shown CPP-CCmut3 exhibits markedly improved internalization and activity against Bcr-Abl compared against wild-type Bcr-CC and untargeted CCmut3^41^. However, translation of the full-length CPP-CCmut3 construct (81 aa) necessitates further design iteration to address the intrinsically high degree of conformational variability, relative to “traditional” small-molecule therapeutics, that predisposed it to loss of helicity and proteolytic degradation in serum and intracellularly.

**Figure 1:**
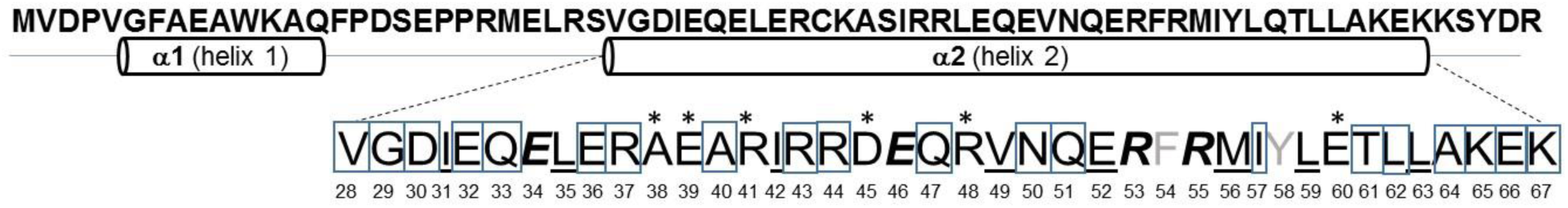
<u>Top sequence</u>, full length cc of Bcr-Abl (helices 1-2); also referred to as Bcr-cc. <u>Bottom sequence</u>, (α2) with rationally designed C38A, K39E, S41R, L45D, E48R, Q60E mutations* to improve heterodimerization with wild-type CC. <u>Underlined black = hydrophobic</u>, ***bold italic = salt bridges***, and <u>underlined gray = interactions with helix 2</u> indicate additional residues that should not be stapled since they are involved in oligomerization. Boxed = possible staple locations. Predicted allowable ***i,i+7* staple locations are: 29&36; 30&37; 33&40; 36&43; 37&44; 40&47; 43&50; 44&51; 50&57; 57&64**.

**Figure 2:**
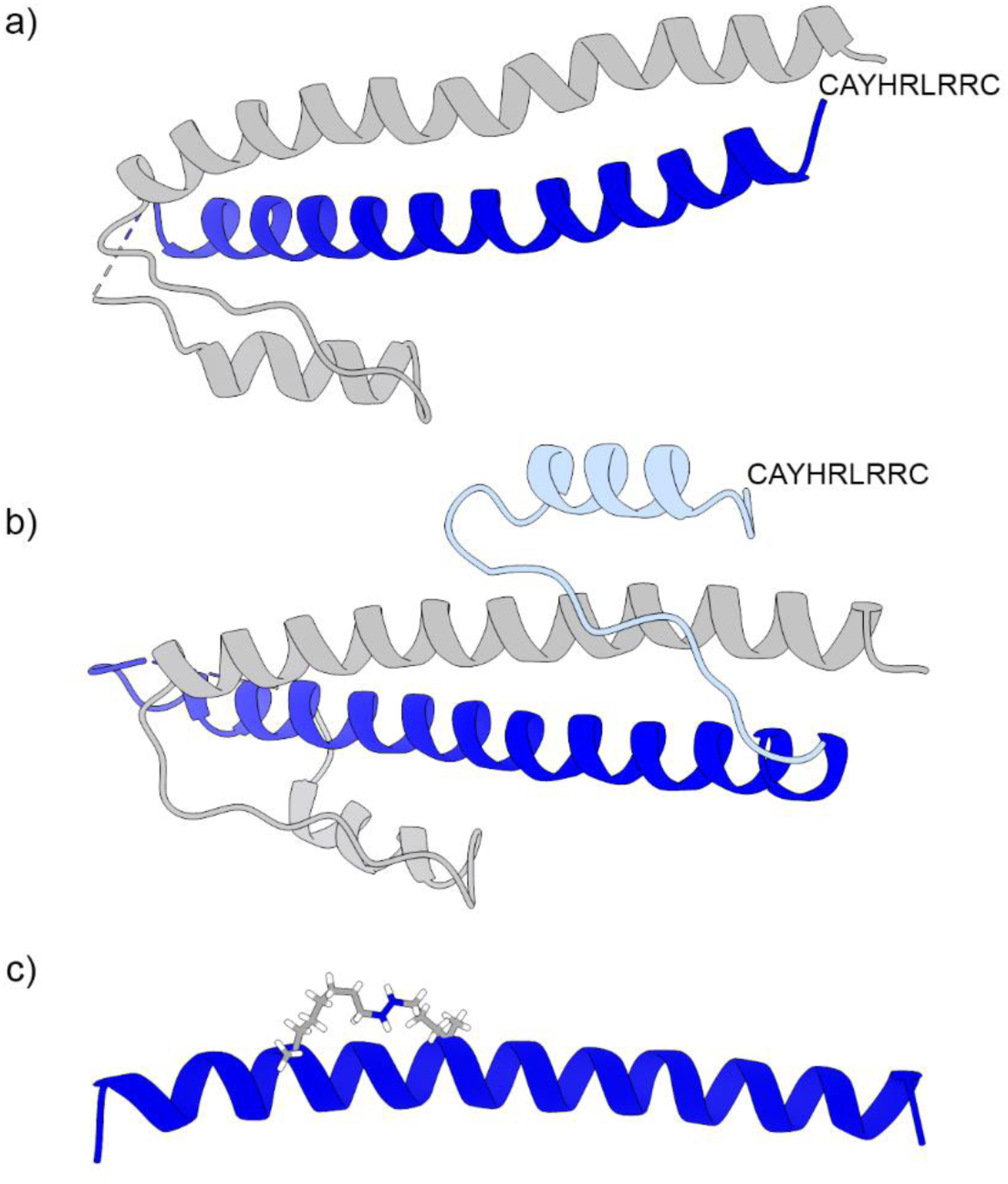
**a)** Bcr-CC:CCmut3-CPP (leukemia-specific cell-penetrating peptide CAYHRLRRC) fused to the coiled-coil domain with C38A, K39E, E48R, S41R, L45D, Q60E mutations in α2 helix (blue) and Bcr-CC domain. **b)** Bcr-CC:CCmut3-CPP (leukemia-specific cell-penetrating peptide CAYHRLRRC) fused to the coiled-coil domain with C38A, K39E, E48R, S41R, L45D, Q60E mutations in α2 helix (blue) with α1 (light blue) + α2 (blue) domain in CCmut3 and Bcr-CC domain (gray). **c)** CCmut3 (α2) stapled – the staple is a hydrocarbon, where the double bond is blue in color.

Herein, we iterate a new generation of our inhibitor against Bcr-CC mediated Bcr-Abl assembly that is designed to reduce conformational variability and improve proteolytic stability through incorporation of all-hydrocarbon “staples” spanning *i, i* + 7 positions in helix α2 (CPP-CCmut3-st). Full-length Bcr-Abl CC encompasses α-helix 1 and α-helix 2, as well as flanking amino acids (aa), spanning residues 1-72 (**Figure 1**). Individual aa residues are chosen to undergo modification (stapling) based on their location in the secondary structure of the peptide. These residues should not be involved in interaction with the target, and must exist in spacings around 1 or 2 full helical turns in the peptide; staple placement of *i, i+7* covers ∼2 helical turns in the protein, providing maximal helical stability/protease protection; thus our initial modeling will be done with *i, i+7* staples (single and double). Examples of possible “ideal” *i, i+7* staple locations (used individually as single staples, or as a double staple) include G29&E36 and N50&I57, which are on the backside of the helix and not in the dimerization interface surface. A comprehensive set of staple sites is shown in **Tables 1-2**. In one study, while single staples exhibited 6-8-fold longer half-lives (vs. unstapled), double staples had a 24-fold enhancement in protease resistance^42^. Using all-atom MD simulations, the Cheatham group previously investigated the structure and dynamics of a subset of *i, i+7* single and double staple locations with a truncated CCmut3, informing an additional staple location (F54&T61), <u>not</u> predicted “manually.” These 11 staple locations were modeled as individual single hydrocarbon staples (SHC) or in combination as double hydrocarbon staples (DHC). In summary, molecular staple crosslinks have emerged as an effective strategy to reinforce α-helices and stabilize peptides by constraining their helical conformation using unnatural α,α-disubstituted amino acids^43^. Moreover, all-hydrocarbon staples have been widely reported to confer proteolytic resistance, enhance serum stability and half-life, and improve cellular uptake^43–52^.

**Table 1:** Systems without staples (used to build the systems with staples) in each case below.

| System without Stapled | Heterodimers |
| --- | --- |
| CCmut3 truncated (CCmut3T or CCmut3) | 111 residues |
| CCmut3 truncated with CPP (CCmut3TC) | 121 residues |
| CCmut3 truncated with CPP (Open) (CCmut3TC') | 121 residues |
| CCmut3 full-length (CCmut3F) | 143 residues |
| CCmut3 full-length with CPP (CCmut3FC) | 153 residues |
| CCmut3 full-length with CPP (Open) (CCmut3FC') | 153 residues |

\*for single stapled systems, the CPP part was tested in cyclic and opened forms

**Table 2:**
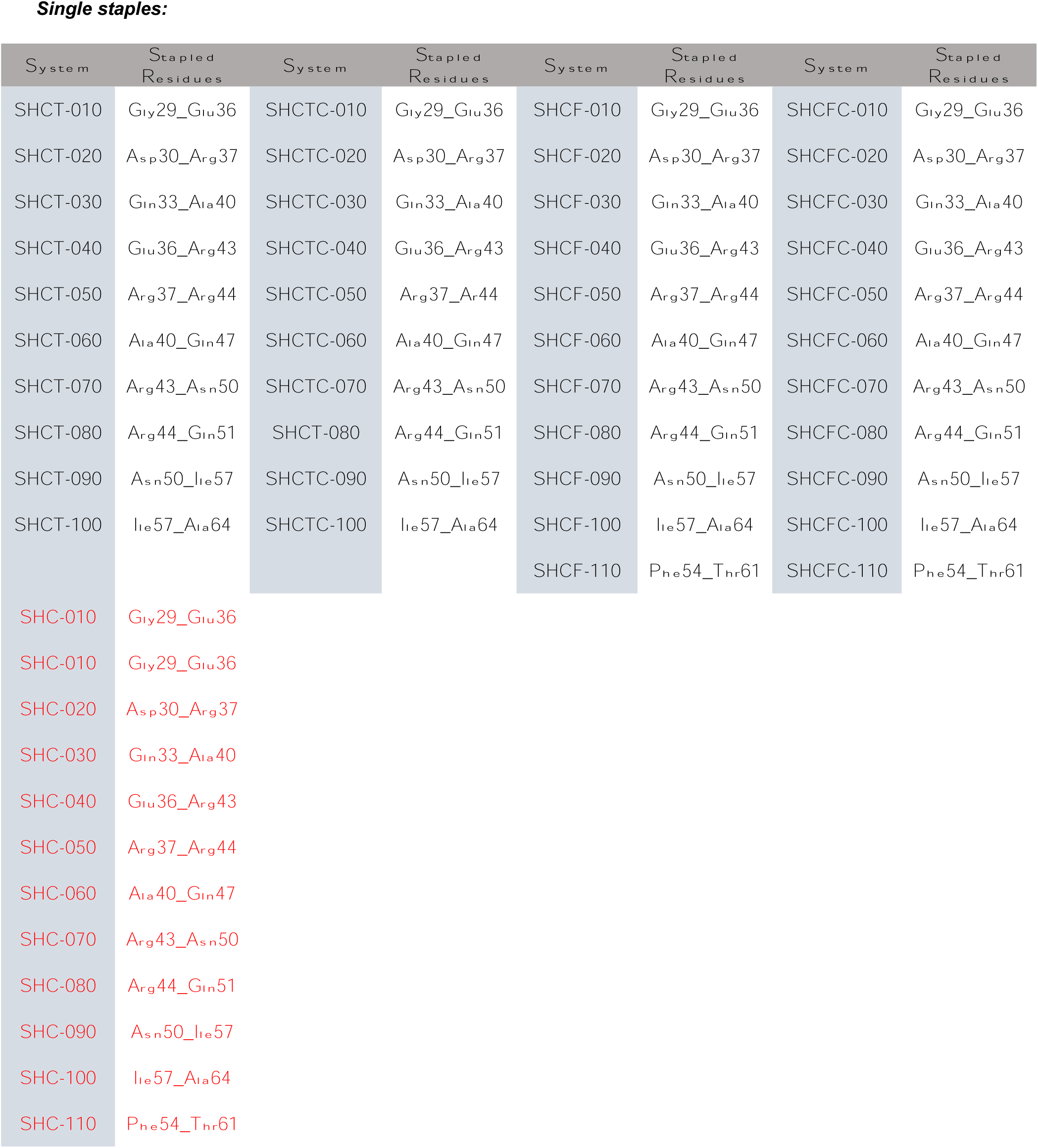
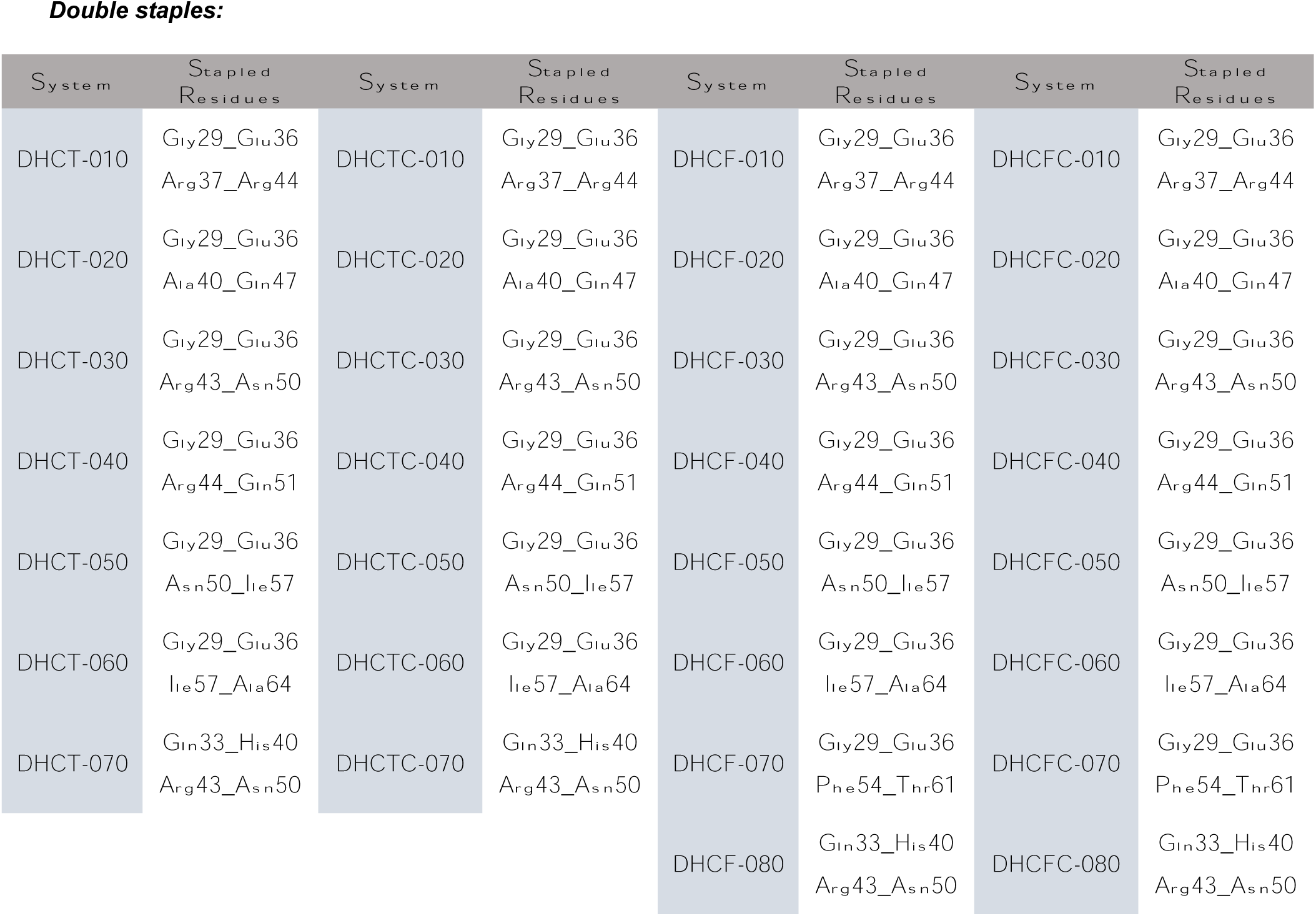
Complete nomenclature for all systems to be modeled, with substituted amino acid residues.

The affinity and specificity of PPIs are intrinsically sequence-specific; thus caution must be exercised in the selection of staple sites such that substitution does not meaningfully diminish target affinity or alter the mechanism of binding^38^. Additionally, large constructs such as CCmut3 possess a multitude of potential staple sites and can accommodate numerous staples in a single construct, economically and temporally precluding the possibility of synthesizing and biophysically characterizing a full library of candidates. Strategies commonplace in stapled peptide design such as sequence-scanning and rational mutagenesis using high-resolution structures are therefore individually insufficient for our interests in their inability to predict behavioral outcomes and screen a reasonably sized library^49,53^. Instead, we utilize computational modeling and biomolecular simulation to design and characterize single and double staple candidates *in silico,* evaluating binding energetics and building upon our seminal work modeling single hydrocarbon staples when applied to a truncated Bcr-CC sequence^38^. This strategy enables us to efficiently build, characterize, and screen lead single/double stapled CPP-CCmut3-st candidates for experimental studies and validation *in vitro* and *in vivo*. In addition to full-length CPP-CCmut, we model a truncated system characterized by deletion of helix α1 and the flexible-loop linker, which are known to impart high conformational variability. To study the impact of the N-terminal cyclic CPP toward model stability and inhibitor activity, we also model the full-length and truncated systems without CPP, with cyclized CPP, and with linear CPP, for a total of six systems (**Table 1**). To these systems, we apply single and double staples identified by sequence scanning of helix α2, omitting native and mutant residues critical for dimerization^39^.The advantage of utilizing computational tools in this endeavor is evident, as they enable rapid and efficient candidate exploration and characterization in silico. We aim to identify lead compounds with optimal properties, significantly reducing the time and costs required for experimental studies.

## Methods

### Model Building

Modified Bcr-Abl oligomerization domain models were constructed based on the crystal structure of the N-terminal oligomerization domain of Bcr-Abl (Protein Data Bank (PDB) entry 1K1F)^29^. Specifically, residues 1-67 from chains A and B were utilized to generate the models. To align with the wild-type protein, the swap tool within the Chimera software was employed to convert selenomethionine residues to methionine and to replace residue 38 with cysteine, mirroring the wild-type configuration (Protein Data Bank (PDB) entry 1K1F)^54^. This conversion process was conducted for all models.

To construct the modified CCmut3 constructs, a sequential implementation of mutations C38A, S41R, L45D, E48R, and Q60E was performed on chain A (residues 1-67) using the swap tool within Chimera. On the other hand, chain B (residues 68-134) remained unmodified, except for the aforementioned changes required to revert it to its wild-type form. To achieve the truncated CCmut3 construct (residues 28-67), the CCmut3-α1 domain with high conformational variability (residues 1-27) was removed. To obtain the truncated CCmut3 model with the leukemia-specific cell-penetrating peptide (CPP) component, we attached the CPP, CAYHRLRRC, to the N-terminus of CCmut3 (**Table 1**). Additionally, to construct the full-length CCmut3 model and enhance its length and mobility, we incorporated the CCmut3-α1 domain (residues 1-27). Adding the CCmut3-α1 domain increased the model’s overall length, flexibility, and mobility. To construct the full-length CCmut3 model with the CPP (CAYHRLRRC)^17^, we attached the CPP sequence to the N-terminus of CCmut3. This fusion of the CPP to the CCmut3 structure enables the investigation of its potential effects on the overall behavior and function of the protein. Staples were strategically employed to create single and double-stapled structures within four distinct sequences of amino acids, as detailed in . The CPP part was tested in cyclic and opened form in single-stapled systems (with the closed form being the expected conformation *in vitro/in vivo*)^41^. We utilized the AMBER General Amber Force Field (GAFF)^42^ and Restrained Electrostatic Potential (RESP)^55^ charges to incorporate staples into the molecular structure. Table 2 lists all stapled constructs modeled herein.

To introduce the staple-containing residues into CCmut3 coils, we performed residue replacement within the PDB file. Specifically, the modified residues with the incorporated staples were substituted into their respective positions within the CCmut3 structure. The models were built using the AMBER ff14SB force-field parameters^56^. To simulate the molecular environment accurately, the models were explicitly solvated in a truncated octahedron shape. The solvent system consisted of TIP3P water molecules^57^, and a minimum surrounding buffer of 10 Å was maintained around the solute. Net-neutralizing counterions (Na+/Cl-) were incorporated into the system using the ion parameters developed by Joung and Cheatham^58^ to maintain overall charge neutrality. Additionally, 50 extra Na+/Cl-atoms were included to achieve a biologically relevant ion concentration of approximately 200 mM. To enhance the efficiency of the molecular dynamics (MD) simulations, the hydrogen masses of the solute were repartitioned to a value of 3.024 Da. This mass repartitioning allowed for doubling the MD time integration step, enabling a time step of 4 fs^59^.

### Minimization and Equilibration

All steps were performed using double precision on central processing unit (CPU) codes. The minimization protocol utilized in this study is based on the methodology developed and validated by Roe and Brooks^60^. It involves a sequence of energy minimization steps followed by relaxation stages, which incorporate short molecular dynamics (MD) simulations. The purpose of this protocol is to facilitate the gradual relaxation of the system and facilitate the attainment of a stable conformation. During this protocol, the system’s pressure was controlled using a Monte Carlo barostat^61^, while the temperature was regulated using a Langevin thermostat^62^ with a collision frequency.

The initial step of the minimization protocol involves 1000 steps of steepest descent minimization. During this step, strong positional restraints are applied exclusively to the heavy atoms of the peptides in the system. The restraints are imposed using a force constant of 5.0 kcal/mol Å, with the initial coordinates serving as the reference structure. It is important to note that no additional constraints, such as SHAKE^63^, should be applied during this minimization step.

The second step of the minimization protocol involves a 15 picosecond (ps) molecular dynamics (MD) simulation conducted at constant volume and temperature (NVT) conditions. During this step, a time step of 1 femtosecond (fs) is employed, resulting in 15,000 simulation steps. To initialize the MD simulation, velocities are assigned to the system’s particles based on a Maxwell-Boltzmann distribution, which ensures that the system starts with the desired temperature. During this MD simulation, positional restraints are still applied to the heavy atoms of the peptides, utilizing a force constant of 5.0 kcal/mol Å, with the initial coordinates serving as the reference structure. Additionally, any necessary constraints, such as SHAKE for hydrogen atoms, should be applied to maintain the desired geometry of the system. A weak-coupling thermostat regulates the temperature, and the thermostat’s time constant is set to 0.5 ps. This helps maintain the desired temperature throughout the simulation.

The third step of the minimization protocol involves 1000 steps of steepest descent minimization. During this step, medium positional restraints are applied to the heavy atoms of the peptides in the system. These restraints use a force constant of 2.0 kcal/mol Å, with the initial coordinates as the reference structure. It is important to note that no additional constraints, such as SHAKE, should be applied during this minimization step. The focus is solely on the heavy atoms of the larger molecules, allowing them to relax further while other system components can adjust more freely.

The fourth step of the minimization protocol involves 1000 steps of steepest descent minimization. During this step, weak positional restraints are applied to the heavy atoms of the peptides in the system. These restraints utilize a force constant of 0.1 kcal/mol Å, with the initial coordinates serving as the reference structure. It is important to note that no other constraints, such as SHAKE, should be applied during this minimization step. The focus remains on the heavy atoms of the peptides, allowing them to undergo further relaxation while other system components can adjust more freely.

The fifth step involves 1000 steps of steepest descent minimization without positional restraints. No constraints, such as SHAKE, are applied to the system during this step. The purpose of this step is to allow the system to relax further and reach a more energetically favorable conformation. By removing the positional restraints, all atoms in the system can adjust their positions based on the potential energy landscape.

The sixth step involves a 5 picosecond (ps) molecular dynamics (MD) simulation conducted at constant pressure and temperature (NPT) conditions. During this step, a time step of 1 femtosecond (fs) is used, resulting in a total of 5000 simulation steps. To initiate the MD simulation, velocities are assigned to the system’s particles based on a Maxwell-Boltzmann distribution, ensuring that the system starts with the desired temperature. During this NPT simulation, positional restraints are still applied to the heavy atoms of the peptides. The restraints utilize a force constant of 1.0 kcal/mol Å, with the initial coordinates from the previous step (step 5) serving as the reference structure. Additionally, any necessary constraints, such as SHAKE for hydrogen atoms, should be applied to maintain the desired geometry of the system. A weak-coupling thermostat and/or barostat can regulate the temperature and pressure during the simulation. The time constant for thermostat and barostat should be set to 1.0 ps. These parameters ensure effective regulation of the temperature and pressure throughout the simulation.

The seventh step involves 5 picoseconds (ps) of molecular dynamics simulation in the NPT ensemble. During this step, a time step of 1 femtosecond (fs) is used, resulting in a total of 5000 simulation steps. The initial velocities for this step should be the final velocities obtained from the sixth step, ensuring continuity in the simulation. This allows the system to build upon the equilibrated state achieved in the previous step. Similar to the sixth step, positional restraints are applied to the heavy atoms of the peptides. The restraints employ a force constant of 0.5 kcal/mol Å, with the final coordinates obtained from the fifth step serving as the reference structure. Additionally, any necessary constraints, such as SHAKE for hydrogen atoms, should be applied to maintain the desired geometry of the system. When employing a weak-coupling thermostat and/or barostat to regulate temperature and pressure during the simulation, the time constant for both should be set to 1.0 ps. This helps maintain the desired temperature and pressure throughout the simulation.

The eighth step of the minimization protocol involves an additional 10 picoseconds (ps) of molecular dynamics (MD) simulation in the NPT ensemble. A 1 femtosecond (fs) time step is used, resulting in 10,000 simulation steps. The initial velocities for this step should be the final velocities obtained from step 7, ensuring continuity in the simulation. This allows the system to continue building upon the equilibrated state achieved in the previous steps. During this step, positional restraints are applied to the backbone atoms of peptide residues. This restraint utilizes a force constant of 0.5 kcal/mol Å, with the final coordinates obtained from the fifth step serving as the reference structure. Additionally, any necessary constraints, such as SHAKE for hydrogen atoms, should be applied to maintain the desired geometry of the system. When using a weak-coupling thermostat and/or barostat to regulate temperature and pressure during the simulation, the time constant for both should be set to 1.0 ps. This helps maintain the desired temperature and pressure throughout the simulation, ensuring accurate sampling of the NPT ensemble.

The ninth step involves 10 picoseconds (ps) of molecular dynamics (MD) simulation in the NPT ensemble. A time step of 2 femtoseconds (fs) is used, resulting in 5000 simulation steps. The initial velocities for this step should be the final velocities obtained from step 8, ensuring continuity in the simulation. This allows the system to continue from the equilibrated state achieved in the previous steps. During this step, no positional restraints are applied to the system. The system is allowed to evolve freely without any restrictions on atom movement. However, any necessary constraints, such as SHAKE for hydrogen atoms, should still be applied to maintain the desired geometry and stability of the system. When using a weak-coupling thermostat and/or barostat to regulate temperature and pressure during the simulation, the time constant for both should be set to 1.0 ps. This helps maintain the desired temperature and pressure throughout the simulation, ensuring accurate sampling of the NPT ensemble.

### Production Molecular Dynamics Simulations

All production molecular dynamics (MD) simulations were performed using the graphics processing unit (GPU) implementation of the Amber20 modeling suite^64^. The simulations were run on 1080ti GPUs. The MD simulations were carried out in an explicit solvent, employing a 4 femtosecond (fs) time step. A Langevin thermostat with a collision frequency of 10 ps^-1^ was used to regulate constant temperature and pressure. This thermostat allows the system to equilibrate and maintain the desired temperature throughout the simulation. A nonbonded cutoff of 10 Å was applied to calculate nonbonded interactions within the system. The particle mesh Ewald (PME) method^65^ handled long-range electrostatic interactions, accurately treating electrostatic forces. The SHAKE algorithm was applied to constrain the bond lengths involving hydrogen atoms. Three independent copies of each system were simulated for statistical purposes and to account for variability. A different random seed was used for each simulation to prevent synchronization artifacts. Each simulation for each system in this study was conducted for approximately 5 microseconds (μs) of simulation time.

### Analysis Protocols

Energetic analyses used the molecular mechanics Poisson-Boltzmann surface area (MM-PBSA)^66^ methodology to estimate the relative binding energetics between each stapled peptide (or unstapled CCmut3) and the Bcr-CC receptor. The MM-PBSA method is commonly used to calculate binding free energies in molecular systems. The MM-PBSA.py is available with the AmbleTools23 suite^67^. The analysis of the candidate peptides was executed using the CPPTRAJ toolset, an integral component of the AmberTools23 package. To enhance our understanding of peptide-peptide interactions, clustering analyses were conducted to identify prevalent conformations sampled during the simulations. In our analysis, we employed the DBScan (Density-Based Spatial Clustering of Applications with Noise) algorithm for conducting clustering. This algorithm is well-suited for identifying clusters within complex and noisy datasets^68^. In this study, the analysis of salt bridges within the trajectories was conducted using MDAnalysis^69^. Specifically, we focused on tracking the salt bridge interactions involving critical amino acid residues, namely GLU with its sidechain atoms OE1 and OE2, ASP with OD1 and OD2, and LYS with NZ, as well as ARG with NH1 and NH2. These interactions were characterized by a distance threshold of less than 4.0 Angstroms.

## Results and Discussion

The research involved extensive all-atom molecular dynamics (MD) simulations, focusing on the structural and dynamic characteristics of various variants of the peptide sequences. Specifically, 93 different variants were examined, encompassing truncated and full-length sequences with and without cell-penetrating peptide (CPP) and single and double-stapled variants featuring a hydrocarbon staple. These candidates were modified by incorporating either one hydrocarbon staple between amino acids *i* and *i* + 7, or two hydrocarbon staples at these positions. The strategic placement of these staples was meticulously carried out along the backbone of CCmut3T, CCmut3TC, CCmut3F, and CCmut3FC sequences (**Table 1**). This placement aimed to ensure that the staples did not disrupt the CCmut3: Bcr-CC binding interface and to preserve any crucial binding partners, such as salt bridges or contacts vital in maintaining the dimerization process. By conducting these MD simulations, the study aimed to unravel the intricate interplay between sequence modifications, stapling, and binding interactions, shedding light on the impact of these factors on the overall structure and dynamics of the peptides.

The following systems nomenclature was utilized for the data in graphs and tables:

1. System Type: SHC means it has a single hydrocarbon staple. DHC is a double hydrocarbon staple.
2. T = truncated CC; F = full length CC
3. Configuration of the CPP: C or C’ represents the specific configuration of the cell penetrating peptide (CPP). C = cyclic/closed and C’ = opened
4. Number: a number is assigned to distinguish between different systems of the same type and configuration. The first number (010) represents the first staple location (pairs of residues) listed in **Figure 2** (29&36); another example is 020, the second set of staple locations (30&37). Residue pairs of staples are listed exhaustively in **Table 2** for both single and double staples.

For example, SHCT-010 has a single staple on a truncated sequence with a staple starting at the first possible designed staple location (29&36, from **Figure 2** and **Table 2**), with no CPP. SHCFC’- 010 would be a single staple on a full-length sequence with a staple starting at the first possible designed staple location with an open CPP configuration.

### Structural Flexibility Analysis

To assess the influence of the staples on structural fluctuations, the Root Mean Square Fluctuation (RMSF) of each system is analyzed and then compared with their counterparts lacking staples. This approach offers insights into how the presence of staples affects the flexibility of the peptide structures. The RMSF values are presented in **Figure 3** and **Figure 4** for both single and double-stapled variants to represent these findings visually. By comparing these RMSF profiles between the stapled and non-stapled systems, we can discern any significant differences in structural fluctuations induced by the stapling modifications. This analysis offers a valuable perspective on the impact of staples at various positions along the peptide sequences and their potential role in stabilizing or altering the overall dynamics of the structures.

**Figure 3:**
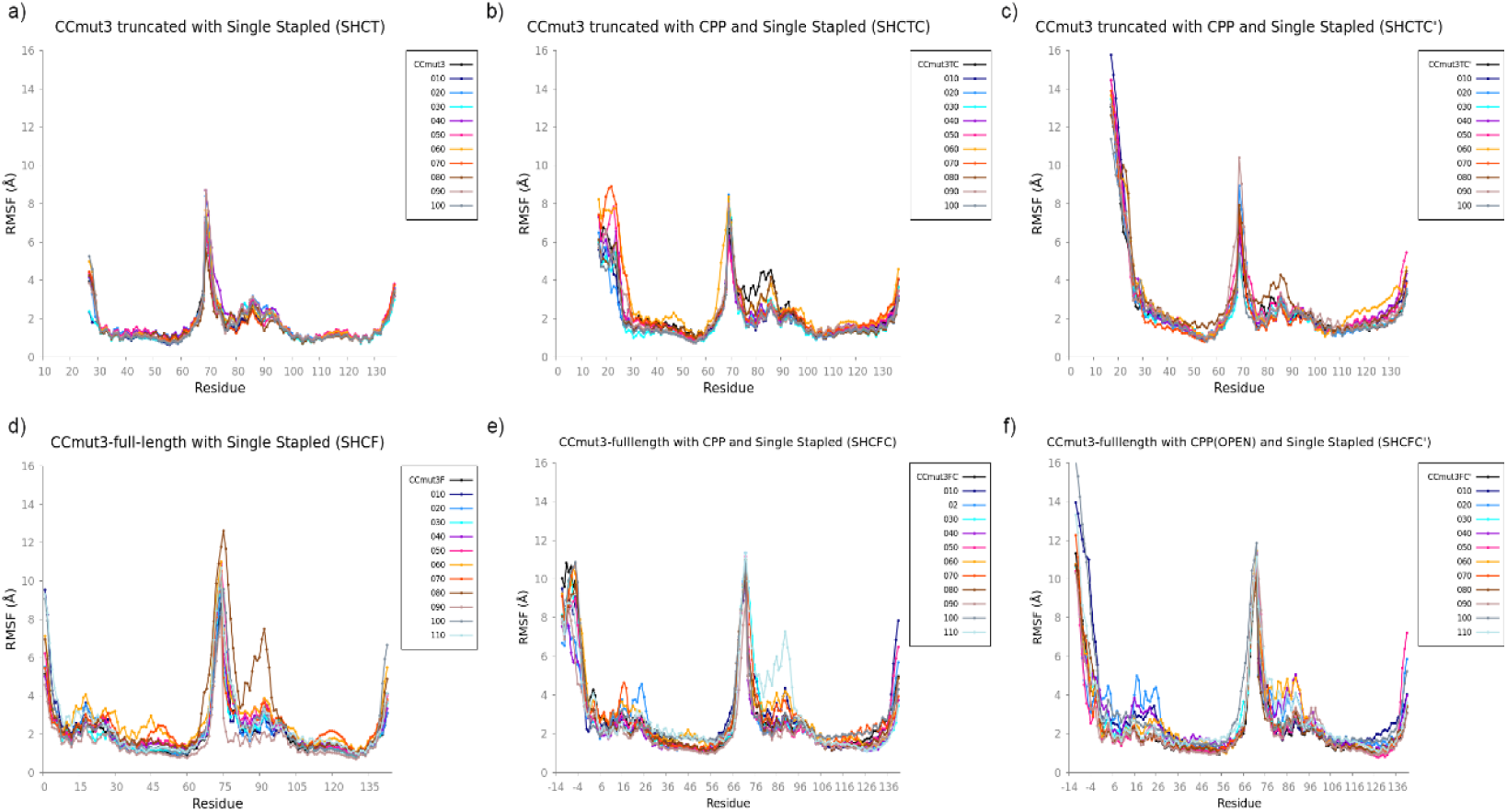
RMSF related to single stapled systems compared to the respective system without staple. **a)** SHCT – CCmut3T with single staple. **b)** SHCTC – CCmut3TC with single staple. **c)** SHTC’ – CCmut3TC’ with single staple. **d)** SHCFC – CCmut3F with single staple. **e)** SHCFC – CCmut3FC with single staple. **f)** SHCFC’ with single staple.

**Figure 4:**
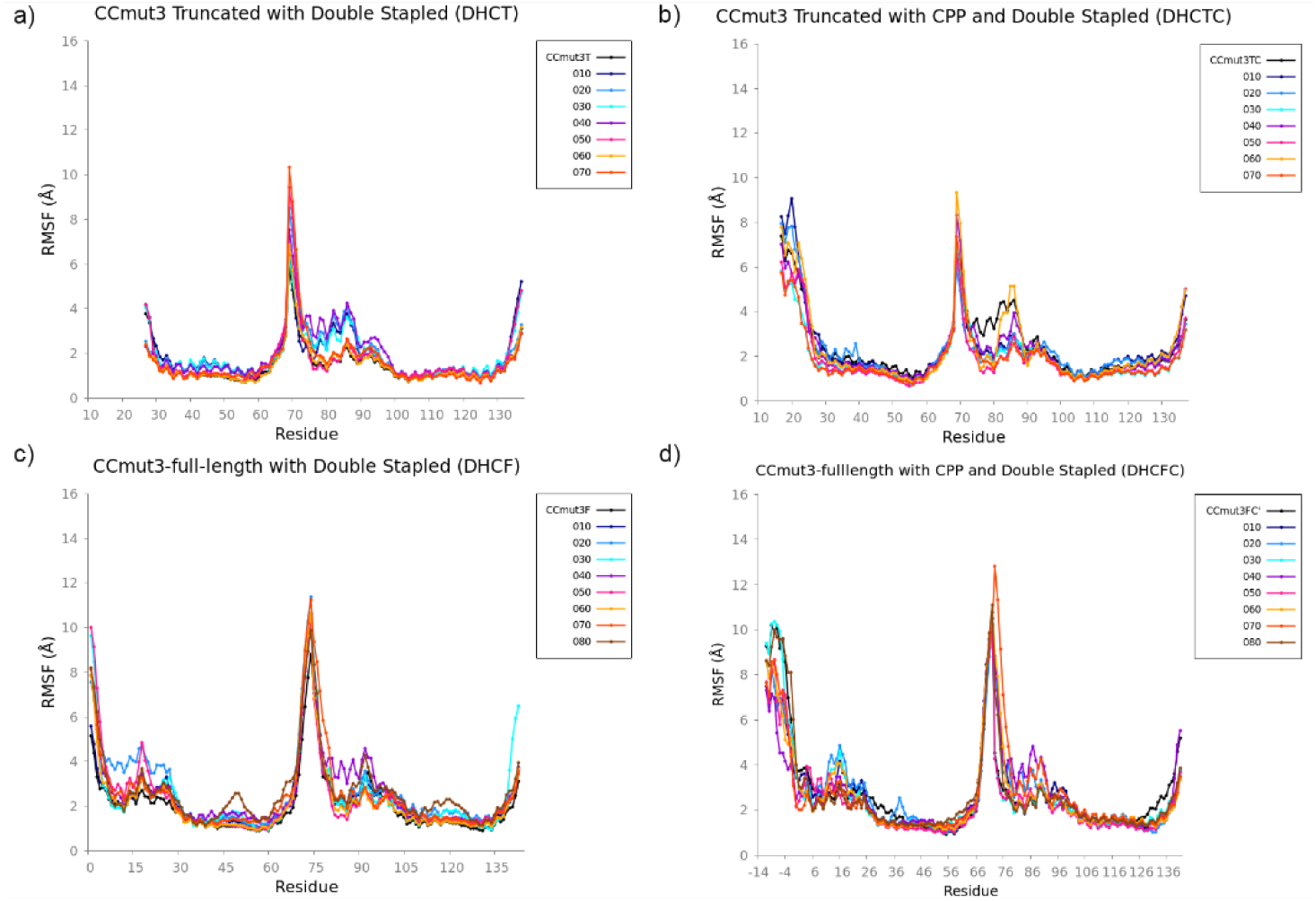
RMSF related to double stapled systems compared to the respective system without staple. **a)** DHCT – CCmut3T with double staple. **b)** DHCTC – CCmut3TC with double staple. **c)** DHCF – CCmut3 full-length with double staple. d) DHCFC – CCmut3FC with double staple.

In the context of systems related to the truncated and single stapled sequence (SHCT), we divided the sequence into two segments: residues 28 to 67, representing CCmut3, and residues 68 to 139, representing Bcr-CC. Our investigation centered on placing a staple along the CCmut3, a key element of interest. As illustrated in **Figure 3a**, our RMSF analysis revealed that the presence of the staple did not induce significant changes in the overall fluctuation compared to the system without the staple. Notably, we observed that the terminal regions exhibited the highest fluctuations. Specifically, the loops and the region connecting α1 and α2 of the wild-type Brc-CC (residues 67-139) demonstrated the greatest flexibility. However, we also observed increased fluctuations in the region corresponding to α1 of Bcr-CC. This region holds particular significance as it is involved in π-stacking interactions between α2 of CCmut3 and α1 of Bcr. Furthermore, our analysis unveiled distinct fluctuations between different SHCT systems. In the SHCT-020 system, we observed a higher degree of fluctuation, indicating increased dynamics in this system. In contrast, the SHCT-070 system displayed comparatively lower fluctuations, suggesting greater stability in this configuration.

Like previous systems, the distinguishing factor in these scenarios is incorporating a cell-penetrating peptide (CPP) at the start of the amino acid sequence. In **Figure 3b**, the CPP is depicted in a closed state, while in **Figure 3c**, the CPP is presented in an open state. There are distinct fluctuation patterns in these two configurations. We observe reduced overall fluctuation in the system where the CPP is open. This is primarily due to α-helices forming within the system, imparting structural stability. However, it is worth noting that part of the structure retains a loop conformation, contributing to increased RMSF value (**Supplementary Information: Helical Analysis**). Conversely, in the case where the CPP is closed, no α-helix formation is observed. This change also leads to variations in the behavior of the rest of the molecular interactions, as evident from the RMSF profiles.

We observe higher fluctuations in the SHCTC-060 system, particularly in the staple region (residues staple 40 and 47). These fluctuations may indicate an ongoing attempt by the system to adapt or adjust to the introduced changes. Such behavior is not observed in the system with the opened CPP. In the Bcr-CC alpha region, the SHCTC-080 and SHCTC’-080 systems exhibit elevated fluctuation values. This suggests distinctive regional interactions or structural features that contribute to the observed fluctuations.

For the RMSF analysis of the full-length sequence, we have provided **Figures 3d** (SHCF), **3e** (SHCFC), and **3f**. In **Figure 3d**, representing the complete sequence without the CPP, we observe increased fluctuation compared to the truncated variant. This is due to the addition of the α1 structure in CCmut3, which leads to heightened interactions between CCmut3 and Bcr-CC, consequently increasing the overall fluctuation. Specifically, we note notable fluctuations in the α1 region of CCmut3 (residues 1-27). This fluctuation is particularly pronounced in the SHCF-060 system. There is significant fluctuation in the α2 regions of the same peptide. In the SHCF-080 system, we observe substantial fluctuation in the α1 region of Bcr-CC, indicating significant movement and dynamics in this specific region. The smallest fluctuation is observed in the SHCF-090 system, suggesting a more stable and well-accommodated state for the system. This reduced fluctuation indicates that the SHCF-090 system may be experiencing a more favorable conformation or interaction, contributing to its enhanced stability.

The results from the SHCFC (**Figure 3e**) and SHCFC’ (**Figure 3f**) systems concerning the region associated with the CPP are consistent with the truncated case. When the CPP is in an open state, it exhibits higher fluctuations. This behavior can be attributed to the formation of a loop between the α-helix regions. **Figure 3e** shows that the region corresponding to α1 in CCmut3 experiences the most significant fluctuations in the SHCFC-020 and SHCFC-070 systems. Similarly, the region from 116 to 130 in α2 of Bcr-CC also exhibits substantial fluctuations. This suggests a dynamic interaction between α1 CCmut3 and α2 Bcr-CC, which plays a crucial role in maintaining the stability of the dimer. In **Figure 3f**, the largest fluctuation in the α1 region of CCmut3 is observed in the SHCFC’-020 system. However, we do not observe a similar trend in the α2 regions of Bcr-CC. Interestingly, both the SHCFC-110 and SHCFC’-110 systems display increased fluctuations in the α2 region of Bcr-CC, indicating more movement in this region.

Next, the RMSF data for the double stapled systems were analyzed (**Figure 4**). **Figure 4a** shows that the systems DHCT-060 and DHCT-070 display the least fluctuation among all the configurations. In contrast, the remaining systems exhibit higher fluctuation in the region corresponding to α1 of Bcr-CC, suggesting increased mobility in this region. Consistent with our observations in other cases, the terminal regions maintain a loop conformation, and the region spanning residues 68-75 shows a consistent peak in fluctuation. We observe varying fluctuation patterns by focusing on the systems with a closed (cyclic) CPP, **Figure 4b**. Among these, the system with the lowest fluctuation is DHCTC-070, while the control and DHCTC-060 systems exhibit the highest fluctuation levels. Notably, the most substantial fluctuations are observed in the alpha1 region of Bcr-CC, which is responsible for interactions with alpha2 of Bcr-CC.

For the next RMSF analysis, our attention shifts to the full-sequence and double-stapled systems. **Figure 4c** shows that DHCF-070 and DHCF-060 systems consistently exhibit the lowest overall fluctuation. In contrast, DHCF-020, DHCF-050, DHCF-080, and DHCF-040 systems display the highest fluctuation levels in specific regions. Notably, the DHCF-080 system stands out, with the greatest fluctuation occurring in the region where the staples are placed. This suggests increased movement and potential challenges in accommodation in this region. Furthermore, elevated fluctuation is also evident in both the α1 and α2 regions of Bcr-CC.

In **Figure 4d**, we examine the system with the complete sequence and a cyclic CPP, observing similar trends in CPP fluctuation patterns as seen in other systems with a comparable CPP structure. Remarkably, the DHCFC-080 system stands out with the lowest overall fluctuation. While other systems are displaying lower fluctuations in specific regions but higher ones in others, making it challenging to assess their overall impact, the DHCFC-020 system is noteworthy for exhibiting greater fluctuation in the region of its first staple, as well as increased fluctuation in the α1 region of CCmut3.

In summary, our RMSF analysis of various systems, including those with truncated sequences, cyclic CPPs, full-length sequences, and double staples, has provided valuable insights into the fluctuations and dynamics of these molecular configurations. We have observed that cyclic CPPs tend to influence fluctuation patterns, with closed CPPs often leading to reduced overall fluctuations. Additionally, we noted variations in fluctuation levels in specific regions of interest, particularly those involving staple placement and crucial interaction sites within the biomolecules.

### Binding Affinity and Thermodynamic Insights

The analysis of ΔG values, representing the binding free energy of various systems, plays a pivotal role in understanding molecular interactions and their thermodynamic aspects. These values serve as a crucial metric in evaluating the stability and affinity of molecular complexes. Our study comprehensively analyzed ΔG values associated with different molecular systems and conditions. Our primary objective is to explore potential relationships within the data, shedding light on the impact of various system parameters on binding affinity.

Our study surrounds diverse systems, each characterized by distinct factors. These factors may include variations in molecular sequences, truncations of sequences, and the presence or absence of cell-penetrating peptides. By investigating these different parameters, we aim to discern patterns or trends that might underlie the observed ΔG values (**Figures 4, 5**).

As we can see in **Figures 5a-c**, the CPP sequence makes the ΔG values more negative than the SHCT values (truncated). ΔG values provide valuable insights into the thermodynamics of molecular interactions. A negative ΔG typically indicates a favorable interaction, suggesting that the binding process is energetically favored. A more negative ΔG signifies a stronger binding affinity and a greater likelihood of stable complex formation.

**Figure 5:**
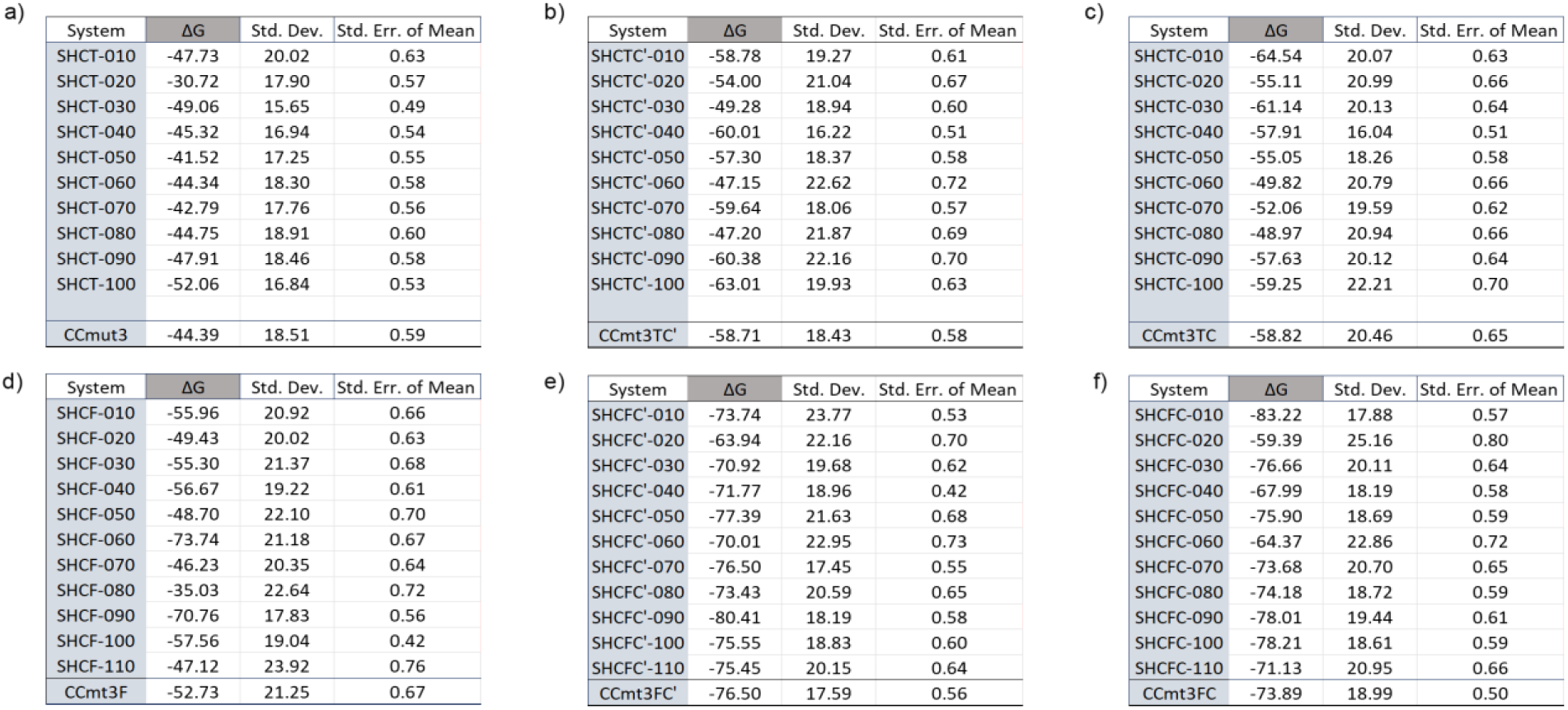
Binding energy for single stapled systems and CCmut3, CCmut3TC and CCmut3F are systems without staples. **b)** SHCTC – CCmut3TC with single staple. **c)** SHTC’ – CCmut3TC’ with single staple. **d)** SHCFC – CCmut3F with single staple. **e)** SHCFC – CCmut3FC with single staple. **f)** SHCFC’ with single staple. Energy unit (kcal/mol).

In addition to our analysis of the ΔG values, we have applied Tukey’s HSD test as a post-hoc method (**Table 3**). This test allows us to make precise comparisons between specific pairs of groups, enabling us to identify statistically significant differences among the conditions we have examined. By employing this statistical tool, we can draw more nuanced conclusions about which systems exhibit meaningful variations in their binding affinities.

**Table 3:** Tukey’s Honestly Significant Difference (HSD) Test Results. Group 1 and Group 2: These columns represent the two groups being compared in Tukey’s HSD test.

a)
| Multiple Comparison of Mean - Tukey HSD, FWER=0.05 |  |  |  |  |  |  |
| --- | --- | --- | --- | --- | --- | --- |
| group1 | group2 | meandiff | p-adj | lower | upper | reject |
| SHCT | SHCTC | -11.53 | 0.001 | -17.69 | -5.37 | TRUE |
| SHCT | SHCTC' | -11.06 | 0.001 | -17.22 | -4.89 | TRUE |
| SHCTC | SHCTC' | 0.47 | 0.9 | -5.69 | 6.63 | FALSE |

| Multiple Comparison of Mean - Tukey HSD, FWER=0.05 |  |  |  |  |  |  |
| --- | --- | --- | --- | --- | --- | --- |
| group1 | group2 | meandiff | p-adj | lower | upper | reject |
| SHCF | SHCFC | -18.75 | 0.001 | -27.05 | -10.44 | TRUE |
| SHCF | SHCFC' | -19.33 | 0.001 | -27.63 | -11.02 | TRUE |
| SHCFC | SHCFC' | -0.58 | 0.9 | -8.88 | 7.73 | FALSE |

The test indicates that there are significant differences between at least some of the groups. In this case, all three group comparisons were statistically tested. (Table 3) A significant difference exists between the SHCT group and SHTCC (cyclic) and SHTCC’(open). This is indicated by the “True” in the “reject” column for these comparisons. There is no significant difference between SHTCC (cyclic) and SHTCC’(open). This is indicated by the “False” in the “reject” column for this comparison. The post-hoc Tukey’s HSD test results show a significant difference between the SHCT group and SHTCC (cyclic) and SHTCC’(open). Still, there is no significant difference between SHTCC (cyclic) and SHTCC’(open). This information helps you understand the specific group differences within your data. In other words, the fact that the CPP is open or cyclic has no impact on the values, but the fact that the CPP makes a difference for systems that do not have it.

After conducting the same analysis for the tables from Figure 5 d-f, based on Tukey’s HSD test results, there are significant differences in ΔG values between SHCF and both SHFC (cyclic) and SHFC’(open). However, there is no significant difference in ΔG values between SHFC (cyclic) and SHFC’(open), Table 3b. In the context of these analyses, the CPP may play a significant role in influencing the binding affinity of the studied systems. This is suggested by Tukey’s HSD test results, which observe substantial differences in ΔG values when comparing different groups, and these differences are likely associated with the presence or absence of the CPP.

The energetic results of the double stapled systems show differences between the different types of systems, so the Turkey test was also applied to better evaluate the relationships between them. (**Figure 6**). The Tukey’s Honestly Significant Difference (HSD) test results indicate which group pairs have statistically significant differences in binding affinity (ΔG values) (**Table 4**). For group 1 and group 4 (DHCF and DHCTC), the mean difference is approximately -1.87. The p-adj is relatively high (0.9), greater than 0.05. In this case, the confidence interval for the mean difference includes zero. This means that there is no statistically significant difference between DHCF and DHCTC. For group 2 and group 3 (DHCFC and DHCT), the mean difference is approximately 31.59, with a low p-adj (0.001). The confidence interval for the mean difference does not include zero, indicating a significant difference in ΔG values between DHCFC and DHCT. For group 2 and group 4 (DHCFC and DHCTC) the mean difference is approximately 17.01, and the p-adj is 0.001, indicating a significant difference. The confidence interval for the mean difference does not include zero, signifying a significant difference in ΔG values between DHCFC and DHCTC. For group 3 and group 4 (DHCT and DHCTC), the mean difference is approximately -14.57, and the -adj is 0.001, indicating a significant difference. The confidence interval for the mean difference does not include zero, implying a significant difference in ΔG values between DHCT and DHCTC.

**Figure 6:**
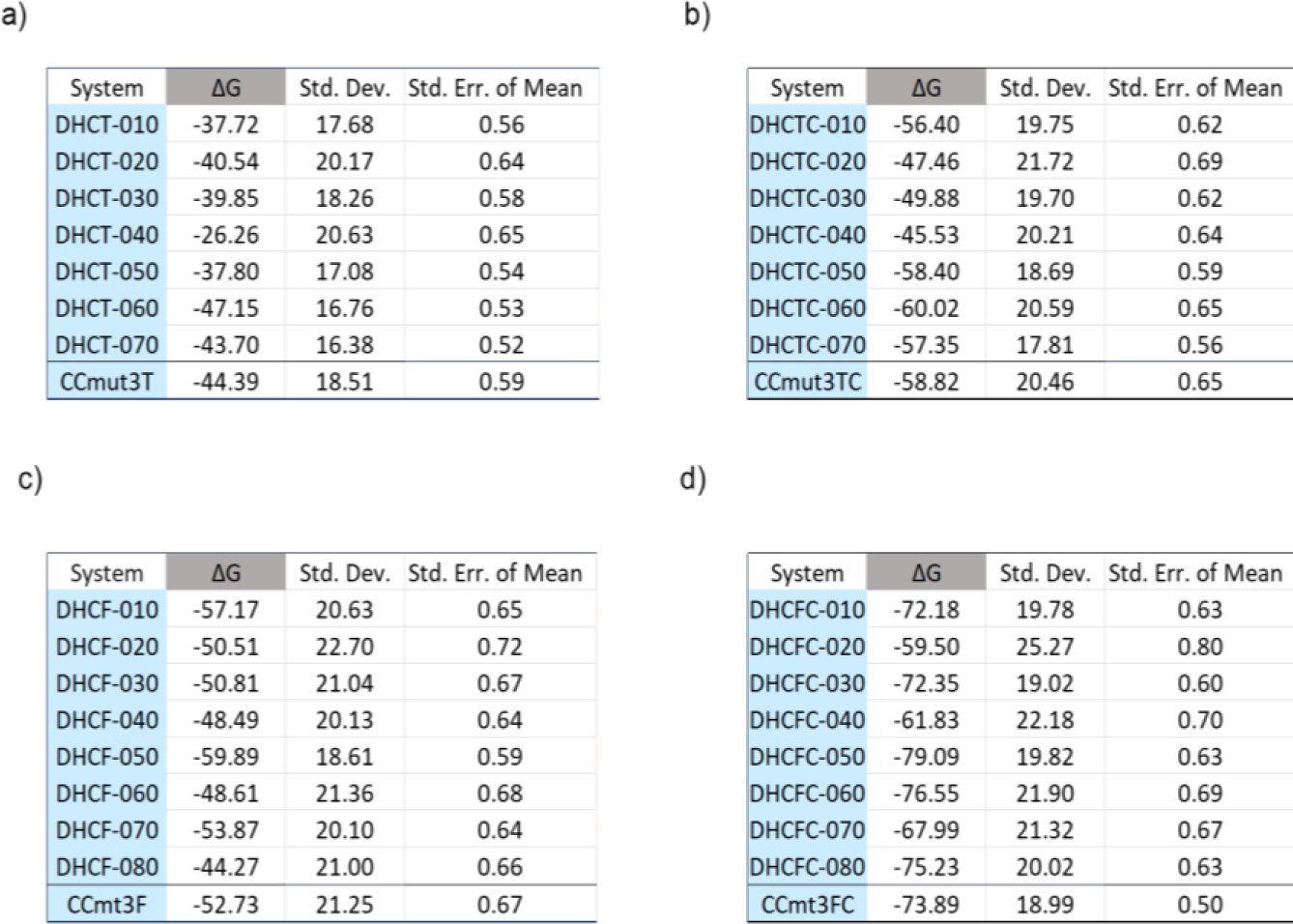
Binding energy for double stapled systems and CCmut3, CCmut3TC and CCmut3F are systems without staples. **a)** DHCT – CCmut3T with double staple. **b)** DHCTC – CCmut3TC with double staple. **c)** DHCFC – CCmut3F with double staple. **d)** DHCFC – CCmut3FC with double staple. Energy unit (kcal/mol).

**Table 4:** Tukey’s Honestly Significant Difference (HSD) Test Results. Group 1 and Group 2: These columns represent the two groups being compared in Tukey’s HSD test.

| Multiple Comparison of Mean - Tukey HSD, FWER=0.05 |  |  |  |  |  |  |
| --- | --- | --- | --- | --- | --- | --- |
| group1 | group2 | meandiff | p-adj | lower | upper | reject |
| DHCF | DHCFC | -18.89 | 0.001 | -27.31 | -10.47 | TRUE |
| DHCF | DHCT | 12.70 | 0.0025 | 3.98 | 21.42 | TRUE |
| DHCF | DHCTC | -1.87 | 0.9 | -10.59 | 6.84 | FALSE |
| DHCFC | DHCT | 31.59 | 0.001 | 22.87 | 40.31 | TRUE |
| DHCFC | DHCTC | 17.01 | 0.001 | 8.30 | 25.73 | TRUE |
| DHCT | DHCTC | -14.57 | 0.001 | -23.58 | -5.57 | TRUE |

**meandiff:** This column indicates the mean difference in $\Delta G$ values between the two groups being compared. **p-adj (Adjusted p-value):** It measures the statistical significance of the mean difference. A low p-adj (e.g., p-adj < 0.05) indicates a significant difference, while a high p-adj suggests that the difference is not statistically significant. **lower, upper (Confidence Interval):** If the confidence interval does not include zero, it indicates a statistically significant difference. If the interval includes zero, it suggests no significant difference. **reject:** The reject column is a binary indicator. If it says "True," the null hypothesis of no significant difference is rejected. If it says "False," it means there is no significant difference between the groups.

Upon analysis of the Tukey’s Honestly Significant Difference (HSD) test results, it becomes evident that there exist substantial disparities in the binding affinity (ΔG values) among various pairs of groups within this study. Specifically, significant differences were observed in the following group comparisons: DHCF vs. DHCFC, DHCF vs. DHCT, DHCFC vs. DHCT, DHCFC vs. DHCTC, and DHCT vs. DHCTC. These findings yield profound insights into the relative binding affinities of these groups. Notably, a compelling trend emerges when we direct our attention to the DHCT and DHCTC cases—comprising the truncated sequence with and without the presence of the cell-penetrating peptide (CPP). Tukey’s HSD test reveals that these two configurations exhibit statistically significant differences in binding affinity. Intriguingly, the energy table depicted in **Figure 5** underpins this observation by demonstrating that the ΔG values associated with DHCTC systems are energetically more favorable when compared to DHCT systems. Likewise, a similar pattern manifests in the DHFC and DHCFC cases. This conveys a crucial message: including the CPP component is a pivotal determinant in the binding affinity of all studied systems. It plays a pivotal role in shaping the thermodynamic landscape of molecular interactions and, by extension, their potential applications.

Incorporating the RMSF data and free energy calculations, we can identify systems within each group that exhibit promising characteristics, such as smaller fluctuations and lower free energy values. For the truncated with single staple (SHTC) cases, the following systems stand out: SHCT-010, SHCT-070, SHCT-100. These systems show both reduced RMSF values, indicating stability and decreased atomic fluctuations, and lower free energy values, suggesting a more favorable energy landscape. Such characteristics would make them noteworthy candidates for further investigation; however, our final therapeutic construct will need to include the closed CPP as this will allow entry into cells. Noteworthy candidates to pursue as therapeutic entities are in bold below (they must contain the closed CPP configuration, designated with C at the end of the name):

Systems with truncated single stapled sequences and the presence of CPP parts are listed as follows:

#### For systems with truncated single staple and CPP

Opened: SHCTC’-010, SHCTC’-030, SHCTC’-040, SHCTC’-070, SHCTC’-090 and SHCTC’-100.

**Closed/Cyclic: SHCTC-010, SHCTC-030, SHCTC-040, SHCTC-070, SHCTC-090, SHCTC-100.**

#### For systems with full-length sequence and single staples

SHCF-010, SHCF-040, SHCF-090 and SHCF-100.

#### For the full-length sequence with single staple and CPP

Opened: SHCFC’-010, SHCFC’-050, SHCFC’-070, SHCFC’-080, SHCFC’-090, SHCFC’-100 and SHCFC’-110.

**Closed/Cyclic: SHCFC-010, SHCFC-030, SHCFC-050, SHCFC-070, SHCFC-080, SHCFC-090, SHCFC-100 and SHCFC-110.**

#### For all the double stabled systems

DHCT-060, DHCT-070, **DHCTC-050, DHCTC-070**, DHCF-010, DHCF-050, DHCF-070, **DHCFC-010, DHCFC-030, DHCFC-050, DHCFC-060, and DHCFC-080.**

These results deepen our understanding of the dynamic behavior of the studied systems and open the door to exciting possibilities in rational design and engineering of biomolecular configurations. The combination of energy profiles and RMSF data has the potential to guide the selection and optimization of systems with desirable stability and dynamics for various applications.

### Impact of Electrostatic Interactions

Our investigation into the formation of salt bridges involving key amino acid residues, such as GLU, ASP, LYS, and ARG, has uncovered crucial insights into the underlying interactions governing the stability and binding affinity of peptide-protein complexes. Salt bridges, facilitated by the electrostatic interactions between oppositely charged residues, can significantly contribute to the overall conformational dynamics and complex stability. Our analysis of these specific salt bridge-forming residues has revealed how their interaction networks evolved over the simulations, shedding light on their role in modulating the structural integrity of the binding interfaces.

To further illuminate the intricate interplay between salt bridge configurations and the stability of peptide-protein complexes, we conducted a cluster analysis to capture representative conformations that feature notable salt bridge interactions (**Figures 7-8**). This approach allowed us to identify and characterize specific salt bridge configurations that could be pivotal in shaping the complex’s structural dynamics and binding affinity. Highlighting specific examples illustrates the impact of salt bridge interactions in the SHCF system and its variants and explains the substantial binding energy differences. To illustrate how salt bridge interactions can underlie significant differences in binding energies, we focus on the SHCF system, a single-stapled full-length variant. Among SHCF-060, SHCF-080 (**Figure 5e**), and CCmut3F (non-stapled) systems, we observe pronounced disparities in their respective binding energy values. Notably, these differences can range on the order of 10 kcal/mol, suggesting the presence of influential molecular interactions that contribute to the observed variations. Upon closer examination of these systems, it becomes apparent that alterations in the salt bridge configurations could provide a plausible explanation for such substantial differences in binding energies. In particular, the presence or absence of stable salt bridge interactions can profoundly impact the overall complex stability and binding affinity. In scenarios where specific salt bridge interactions are established or disrupted, the resulting changes in the electrostatic contributions can lead to marked shifts in binding energy values.

**Figure 7:**
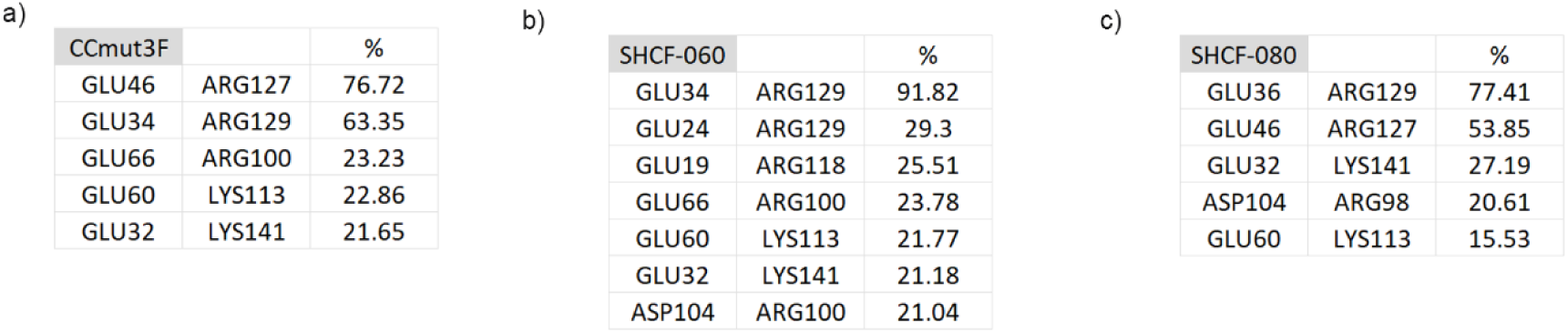
Salt Bridge Analysis – Population of the frames during the simulation that counts the iteration between GLU@OE1, OE2, ASP@OD1, OD2, and LYS@NZ, ARG@NH1, NH2. These interactions are between peptides. **a)** System CCmut3 full-length without staple. **b)** System CCmut3F with single staple position 060. **c)** System CCmut3F with single staple position 080.

**Figure 8:**
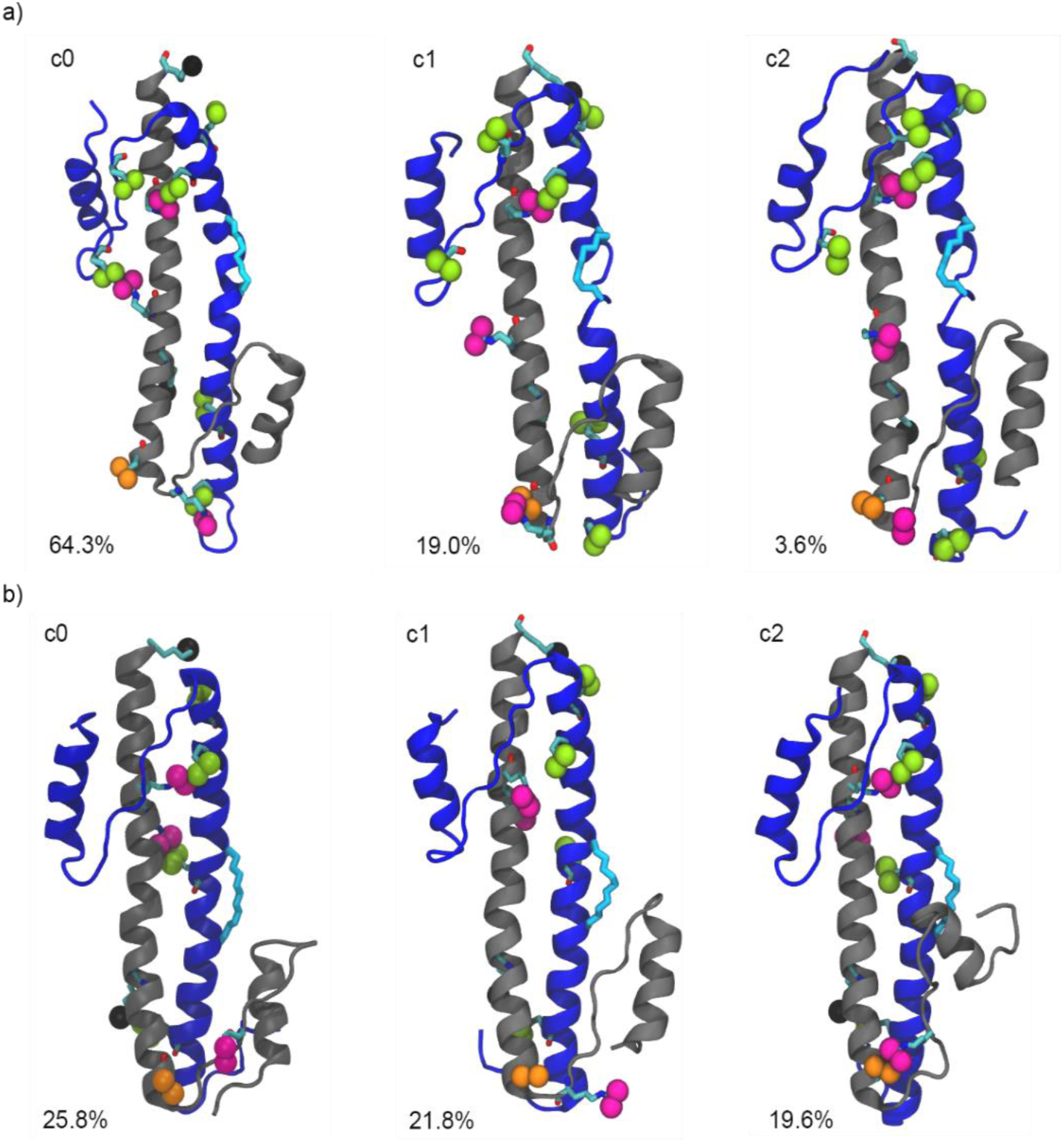
Cluster analysis – Systems a) SHCF – 060. **b)** SHCF – 080. In the images of these structures, highlighting the atoms that participate in forming salt bridges. Specifically, the atoms LYS@NZ are depicted in black, ASP@OD1 and ASP@OD2 are shown in orange, ARG@NH1 and ARG@NH2 are represented in magenta, and GLU@OE1 and GLU@OE2 are denoted in lime. This color-coded visualization effectively delineates the amino acid residues involved in these electrostatic interactions. The helix structure in gray is Br-CC, and blue is the CCmut3F where the staple is placed (licorice-cyan).

Cluster analysis is a powerful tool for elucidating the diversity of structural conformations within molecular dynamics simulations. In our study, cluster analysis has provided a comprehensive snapshot of the most prevalent configurations within the SHCF-060 and SHCF-080 systems. As depicted in **Figure 8**, these representative structures offer a visual insight into the role of salt bridge interactions, further emphasizing the dynamic nature of these critical molecular interactions.

## Conclusions

In these extensive all-atom molecular dynamics (MD) simulation studies, we explored the structural and dynamic characteristics of numerous peptide sequence variants, encompassing truncated and full-length sequences, the presence or absence of cell-penetrating peptides (CPP), and both single and double-stapled configurations. Our meticulous placement of hydrocarbon staples aimed to preserve critical binding interfaces and interactions between CCmut3 and Bcr-CC, shedding light on the intricate interplay between sequence modifications, stapling, and binding interactions. Our nomenclature for the systems allows for clear differentiation and categorization.

### Structural Flexibility Analysis

Our Root Mean Square Fluctuation (RMSF) analysis unveiled critical insights into the influence of staples on structural fluctuations within these peptide systems. While staples did not induce significant changes in overall fluctuations, the terminal regions, particularly loops, and regions connecting α1 and α2 of Bcr-CC, displayed the greatest flexibility. Furthermore, fluctuations in the α1 region of Bcr-CC were observed, vital for π-stacking interactions with α2 of CCmut3. These fluctuation patterns varied between different systems, highlighting the influence of a cyclic or open CPP on structural dynamics.

### Binding Affinity and Thermodynamic Insights

The analysis of ΔG values elucidated the binding free energy, revealing intriguing thermodynamic aspects of molecular interactions. The presence of CPP, whether open or cyclic, had a substantial impact on ΔG values, indicating the significant role of the CPP component in shaping binding affinity. The Tukey’s Honestly Significant Difference (HSD) test further reinforced these findings, showing statistically significant differences in ΔG values for various system configurations. Importantly, this data can guide the selection of promising systems with reduced fluctuations and lower free energy values.

### Impact of Electrostatic Interactions

Our examination of salt bridge interactions involving key amino acid residues provided crucial insights into peptide-protein complexes’ stability and binding affinity. Cluster analysis allowed us to identify representative conformations featuring notable salt bridge interactions, shedding light on their role in modulating structural integrity. The dynamic nature of these interactions was further illustrated through visual representations of representative structures within the SHCF-060 and SHCF-080 systems.

### Lead CPP-CCmut3-st Candidates

Moving forward with experimental proteins to synthesize, we will choose those candidates that had favorable computational characteristics along with the presence of the cell penetrating peptide. Interestingly, some truncated sequences came out as viable candidates (which will simplify the chemical synthesis process, as they are shorter in length). The following are candidates for chemical synthesis and further testing:

### For systems with truncated single staple and CPP

**SHCTC-010, SHCTC-030, SHCTC-040, SHCTC-070, SHCTC-090, SHCTC-100**.

### For the full-length sequence with single staple and CPP

**SHCFC-010, SHCFC-030, SHCFC-050, SHCFC-070, SHCFC-080, SHCFC-090, SHCFC-100 and SHCFC-110**.

### For double stabled systems, both truncated or full length with CPP

**DHCTC-050, DHCTC-070**, **DHCFC-010, DHCFC-030, DHCFC-050, DHCFC-060, and DHCFC-080.**

Although this is still a list of 21 candidates to be synthesized, it is only a subset of all possible candidates. Importantly, this computational study greatly narrows the number of candidates to be synthesized.

In summary, our study not only deepens our understanding of the dynamic behavior of these systems but also opens exciting avenues for the rational design and engineering of biomolecular configurations. The combination of energy profiles, RMSF data, and salt bridge analysis can potentially guide the selection and optimization of systems for specific applications, harnessing the interplay between structure and dynamics. These findings are integral in advancing molecular engineering and biotechnology research.

## Supporting information

Helical Analysis

## Acknowledgement

Research reported in this preprint was supported by the National Cancer Institute of the National Institutes of Health under award number R01CA244583.

