## Supplementary material for "Computational Modeling of Stapled Coiled-Coil Inhibitors Against Bcr-Abl: Toward a Treatment Strategy for CML": Helical Analysis

| | ΔG | Std. Dev. | Std. Err. of Mean | ΔΔG | Std. Dev. | Std. Err. of Mean |
|---|---|---|---|---|---|---|
| DHCT-010 | -37.72 | 17.68 | 0.56 | 6.67 | 25.71 | 0.57 |
| DHCT-020 | -40.54 | 20.17 | 0.64 | 3.85 | 27.48 | 0.61 |
| DHCT-030 | -39.85 | 18.26 | 0.58 | 4.54 | 26.11 | 0.58 |
| DHCT-040 | -26.26 | 20.63 | 0.65 | 18.13 | 27.82 | 0.62 |
| DHCT-050 | -37.80 | 17.08 | 0.54 | 6.59 | 25.30 | 0.57 |
| DHCT-060 | -47.15 | 16.76 | 0.53 | -2.76 | 25.08 | 0.56 |
| DHCT-070 | -43.70 | 16.38 | 0.52 | 0.69 | 24.83 | 0.56 |
| CCmut3 | -44.39 | 18.51 | 0.59 | 0.00 | 26.28 | 0.59 |

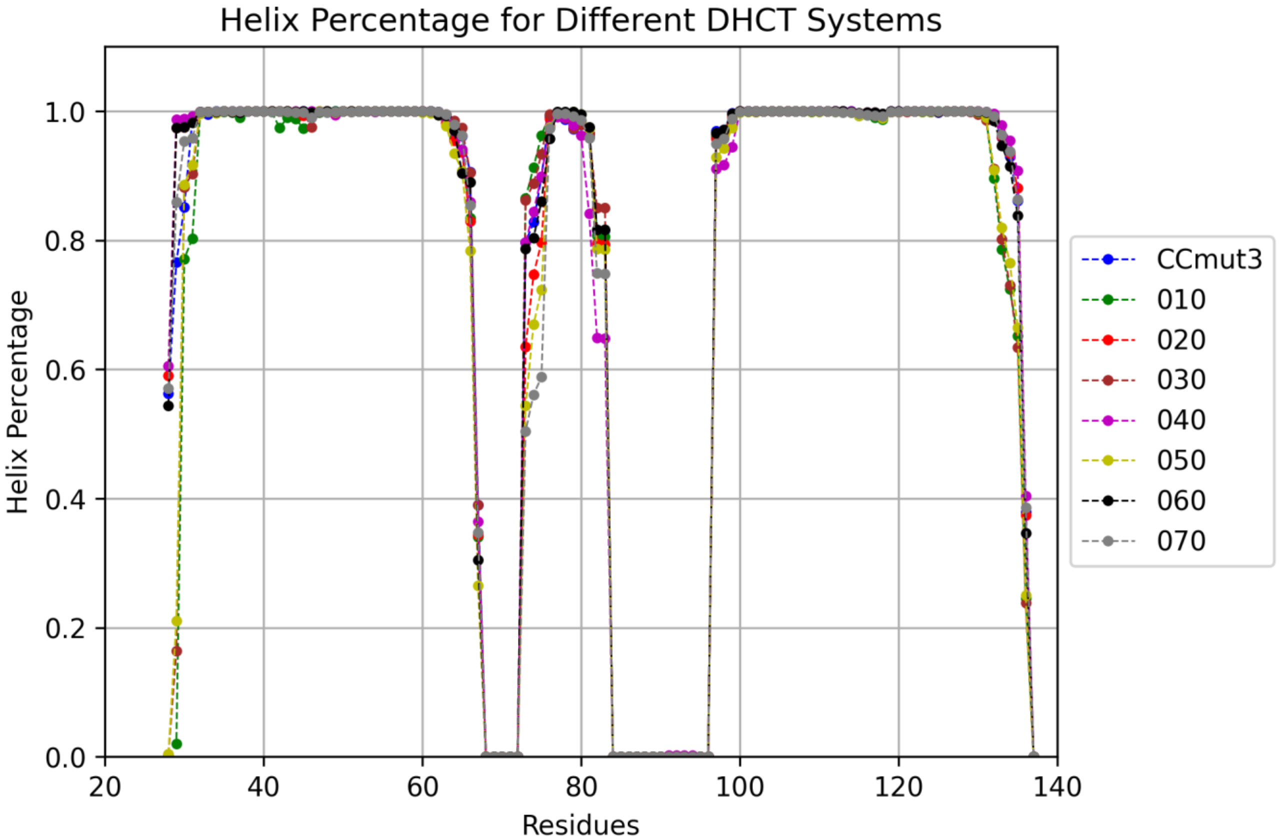


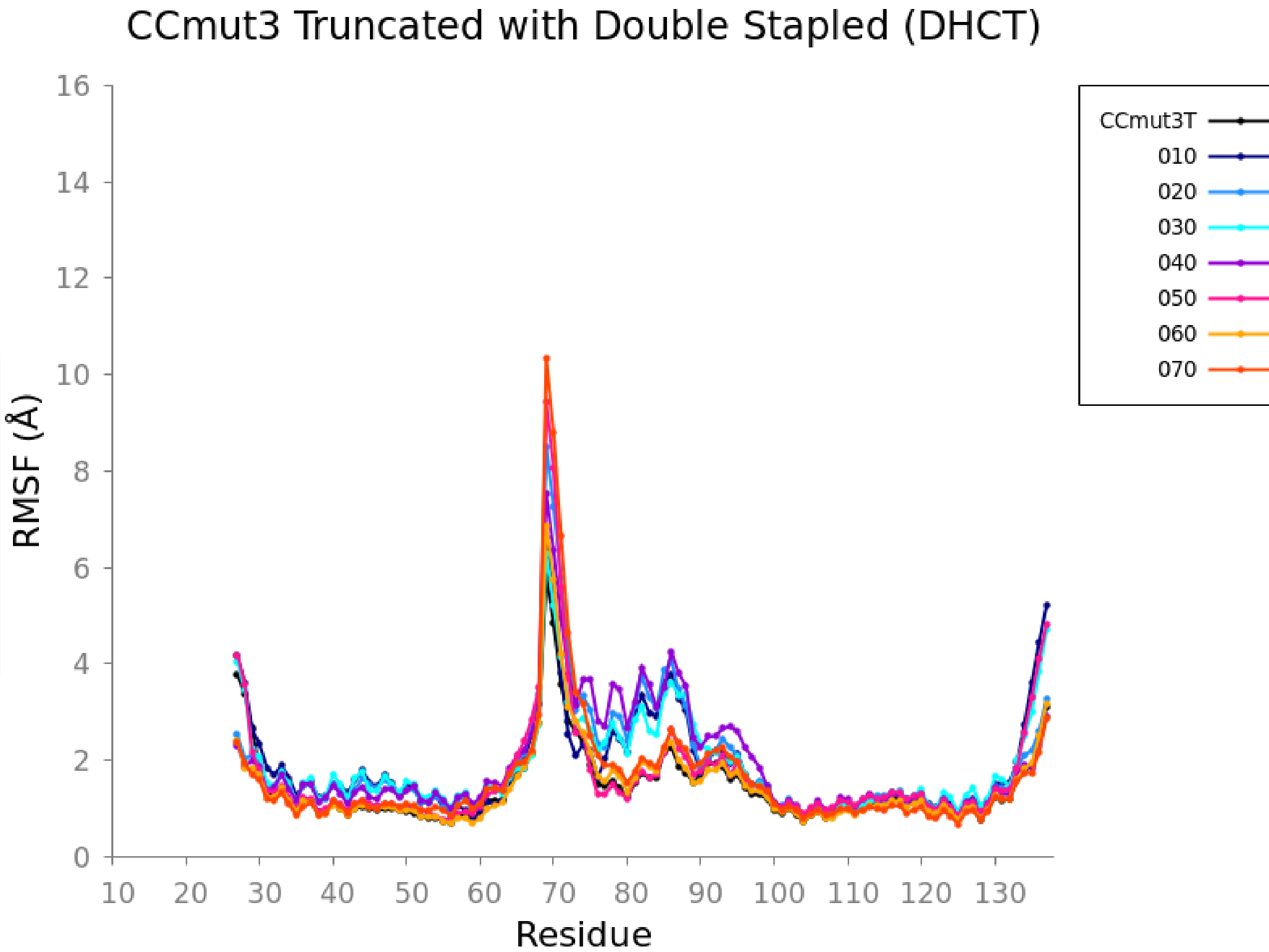

| | ΔG | Std. Dev. | Std. Err. of Mean | ΔΔG | Std. Dev. | Std. Err. of Mean |
|---|---|---|---|---|---|---|
| DHCTC-010 | -56.40 | 19.75 | 0.62 | 2.42 | 28.44 | 0.90 |
| DHCTC-020 | -47.46 | 21.72 | 0.69 | 11.35 | 29.84 | 0.94 |
| DHCTC-030 | -49.88 | 19.70 | 0.62 | 8.94 | 28.40 | 0.90 |
| DHCTC-040 | -45.53 | 20.21 | 0.64 | 13.29 | 28.76 | 0.91 |
| DHCTC-050 | -58.40 | 18.69 | 0.59 | 0.42 | 27.71 | 0.88 |
| DHCTC-060 | -60.02 | 20.59 | 0.65 | -1.21 | 29.03 | 0.92 |
| DHCTC-070 | -57.35 | 17.81 | 0.56 | 1.47 | 27.12 | 0.86 |
| CCmut3TC | -58.82 | 20.46 | 0.65 | 0.00 | 28.93 | 0.91 |

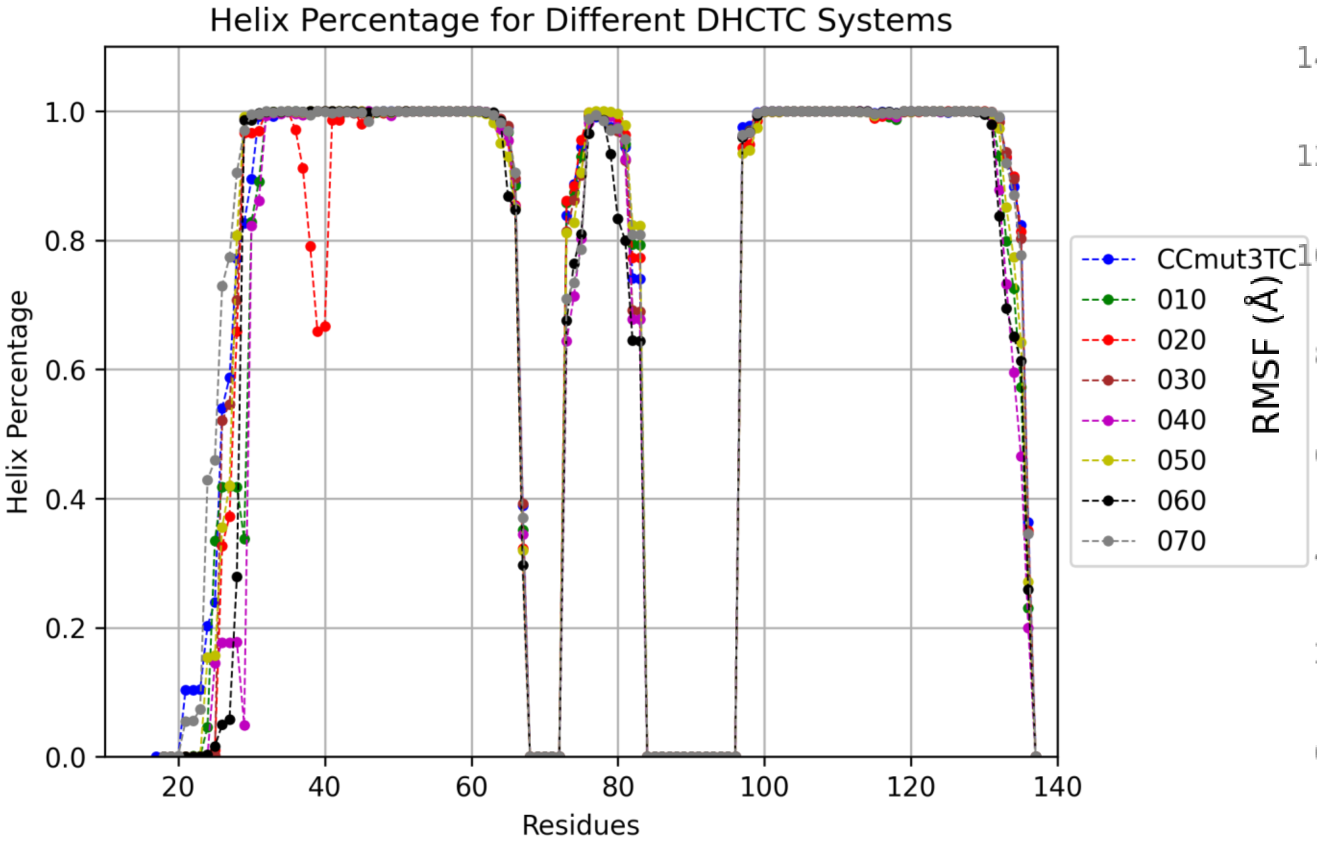


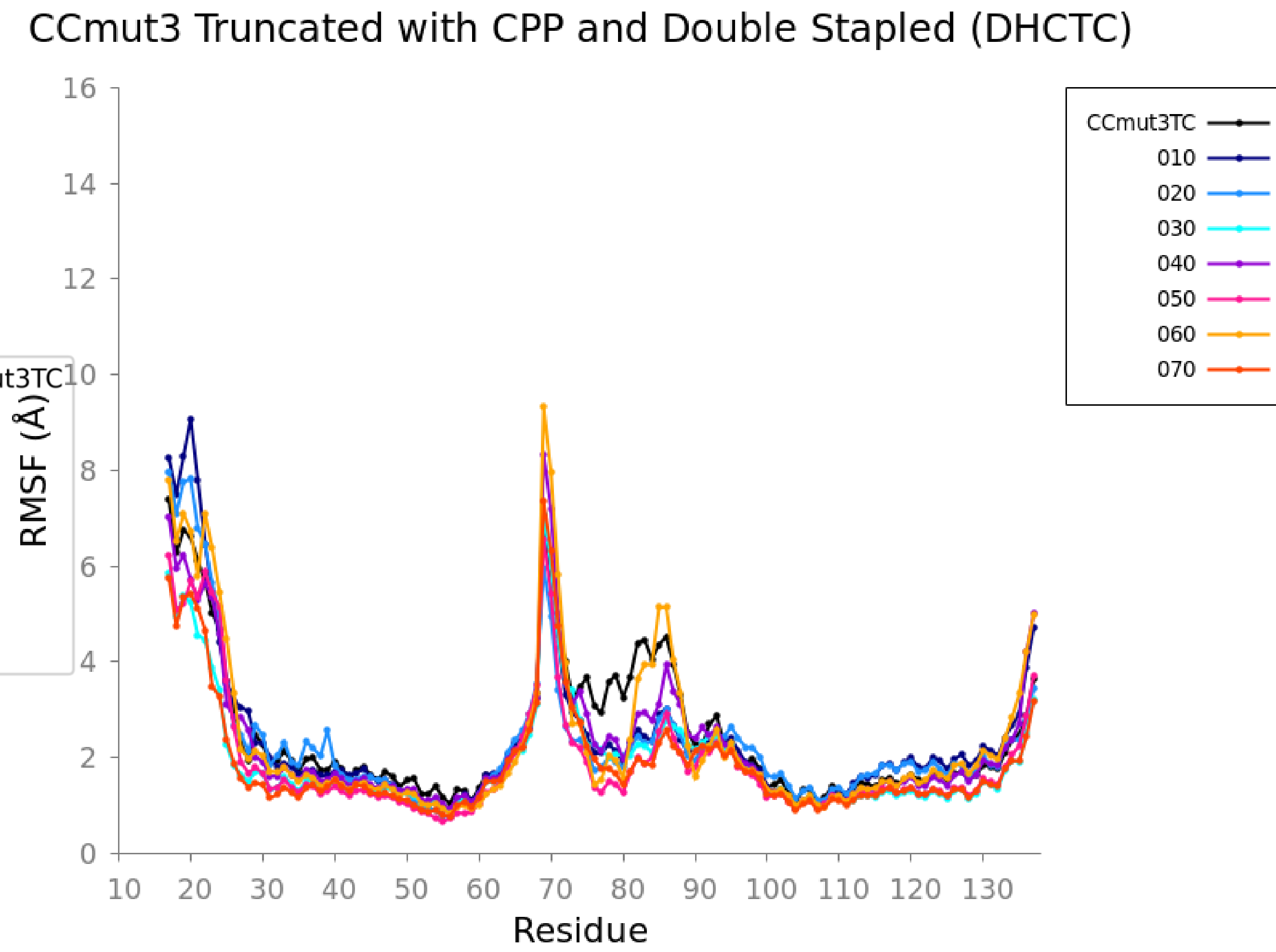

|  | ΔG | Std. Dev. | Std. Err. of Mean | ΔΔG | Std. Dev. | Std. Err. of Mean |
|---|---|---|---|---|---|---|
| DHCF-010 | -57.17 | 20.63 | 0.65 | -4.44 | 29.42 | 0.66 |
| DHCF-020 | -50.51 | 22.70 | 0.72 | 2.23 | 30.91 | 0.69 |
| DHCF-030 | -50.81 | 21.04 | 0.67 | 1.92 | 29.71 | 0.66 |
| DHCF-040 | -48.49 | 20.13 | 0.64 | 4.24 | 29.07 | 0.65 |
| DHCF-050 | -59.89 | 18.61 | 0.59 | -7.16 | 28.04 | 0.63 |
| DHCF-060 | -48.61 | 21.36 | 0.68 | 4.12 | 29.94 | 0.67 |
| DHCF-070 | -53.87 | 20.10 | 0.64 | -1.14 | 29.06 | 0.65 |
| DHCF-080 | -44.27 | 21.00 | 0.66 | 8.46 |  |  |
| CCmt3F | -52.73 | 21.25 | 0.67 | 0.00 | 29.86 | 0.67 |

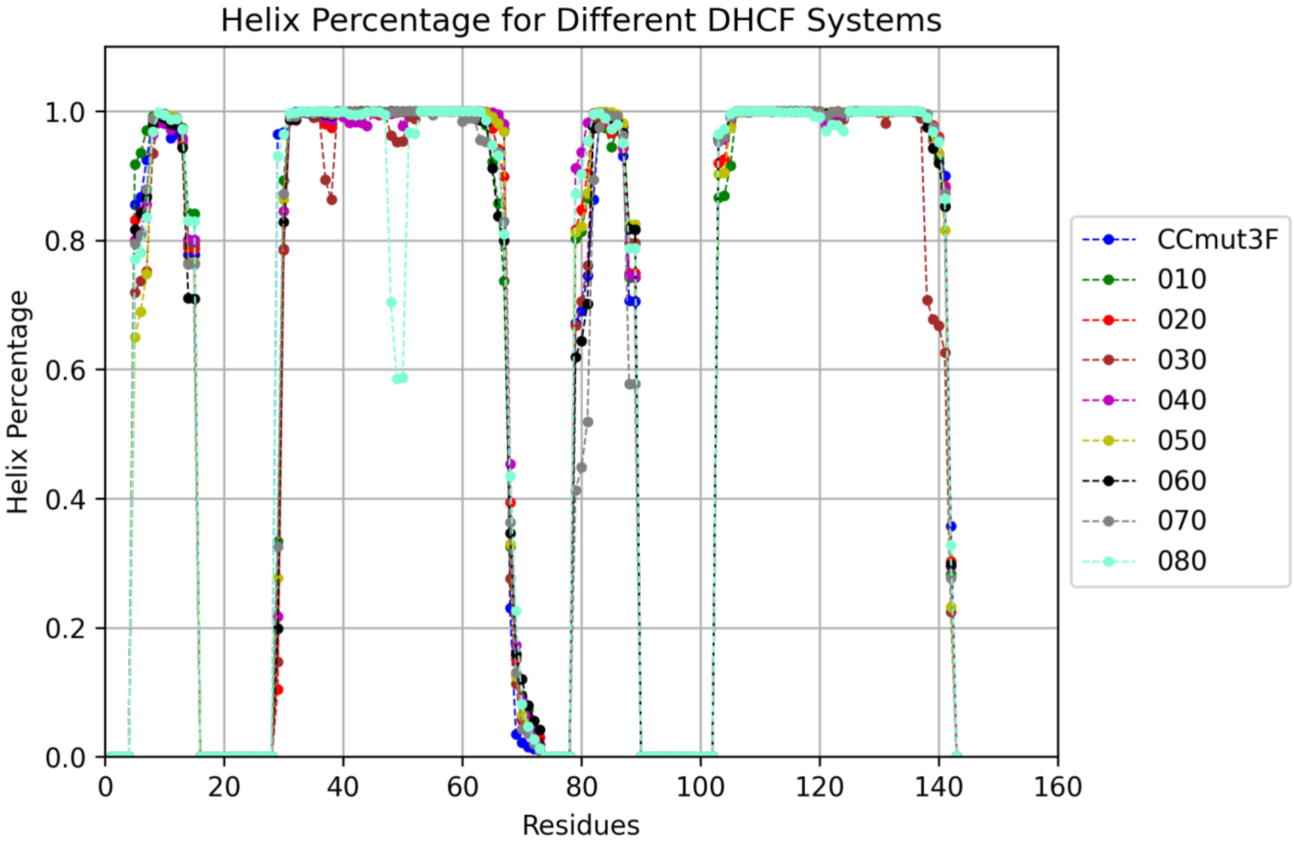


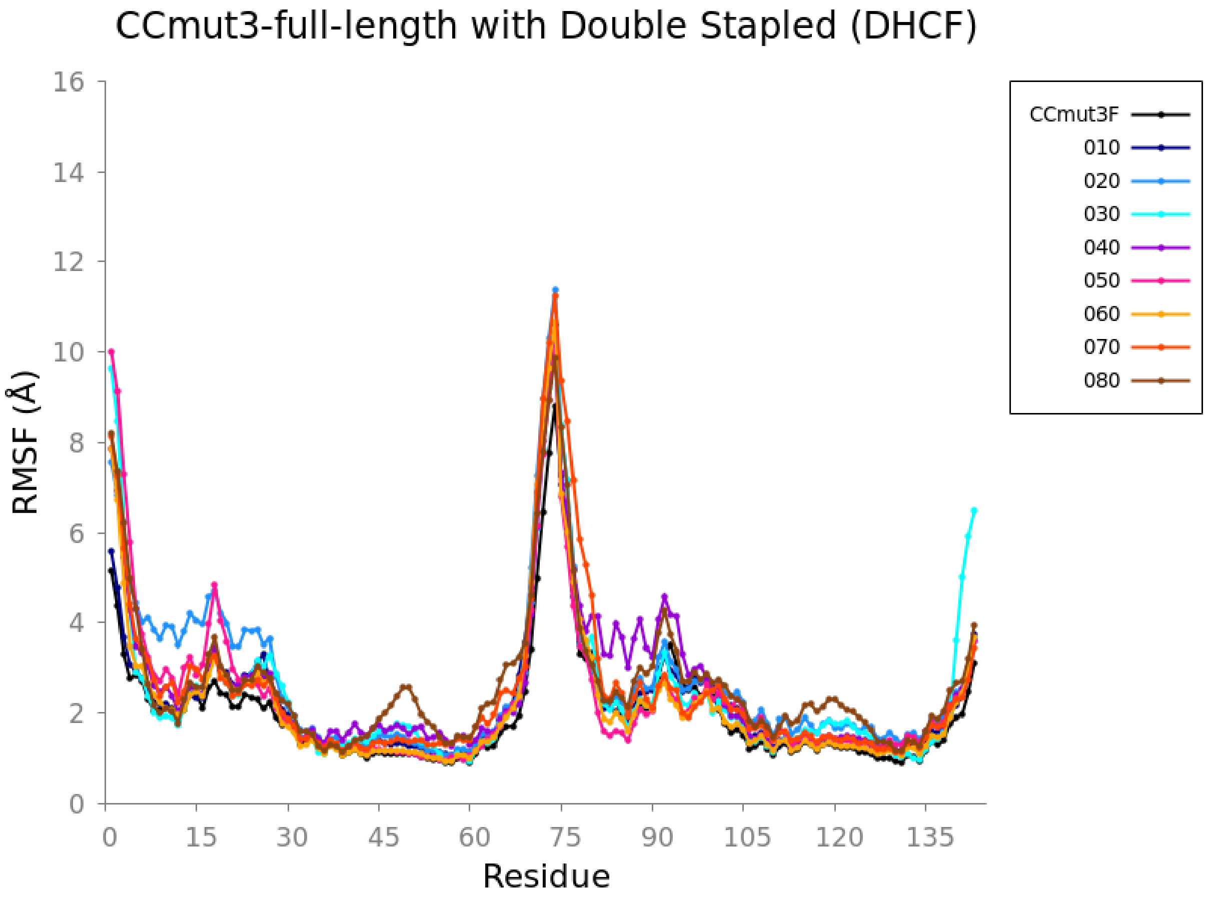

| | ΔG | Std. Dev. | Std. Err. of Mean | ΔΔG | Std. Dev. | Std. Err. of Mean |
|---|---|---|---|---|---|---|
| DHCFC-010 | -72.18 | 19.78 | 0.63 | 1.70 | 27.42 | 0.87 |
| DHCFC-020 | -59.50 | 25.27 | 0.80 | 14.39 | 31.61 | 1.00 |
| DHCFC-030 | -72.35 | 19.02 | 0.60 | 1.53 | 26.88 | 0.85 |
| DHCFC-040 | -61.83 | 22.18 | 0.70 | 12.05 | 29.20 | 0.92 |
| DHCFC-050 | -79.09 | 19.82 | 0.63 | -5.21 | 27.45 | 0.87 |
| DHCFC-060 | -76.55 | 21.90 | 0.69 | -2.67 | 28.98 | 0.92 |
| DHCFC-070 | -67.99 | 21.32 | 0.67 | 5.89 | 28.55 | 0.90 |
| DHCFC-080 | -75.23 | 20.02 | 0.63 | -1.34 | 27.59 | 0.87 |
| CCmt3FC | -73.89 | 18.99 | 0.50 | 0.00 | 26.85 | 0.85 |

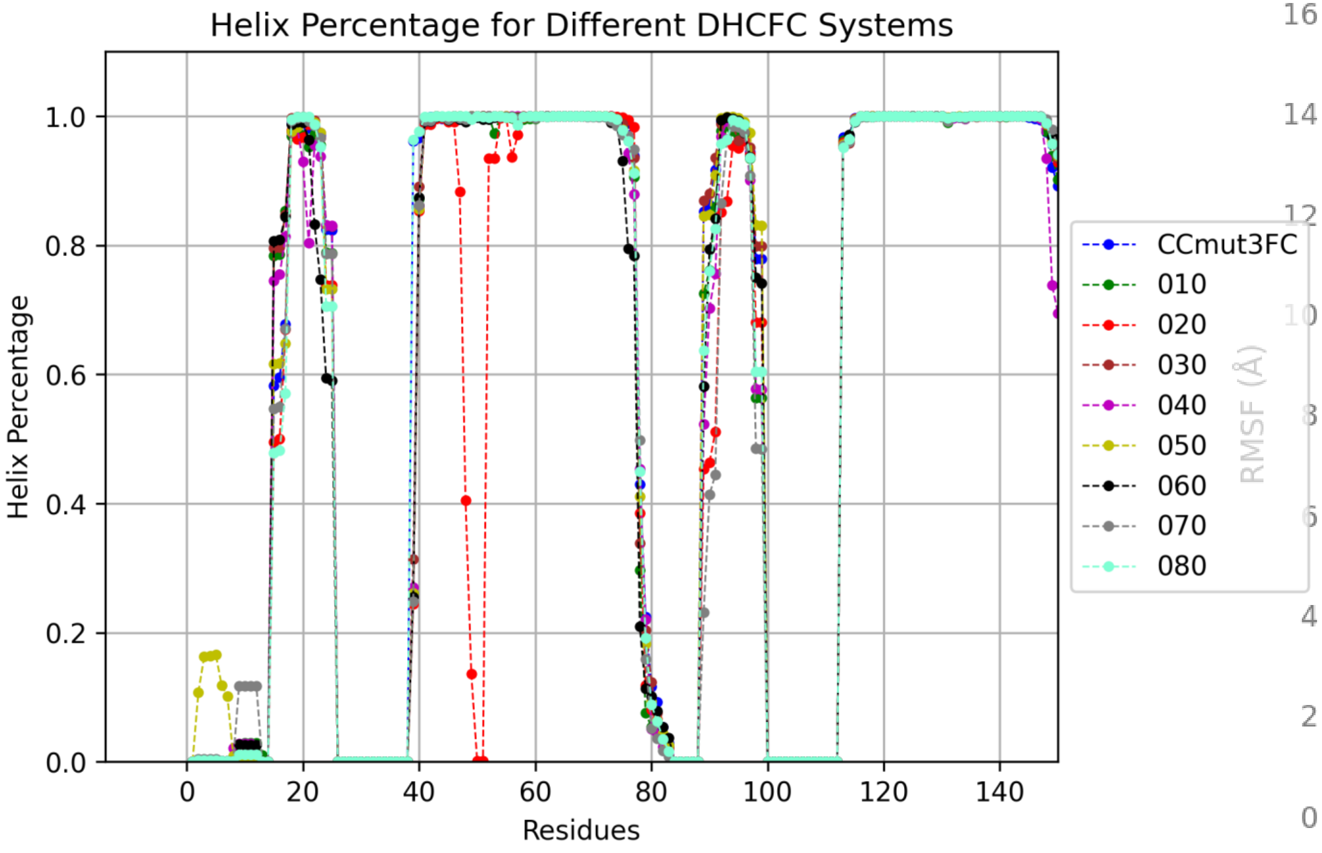


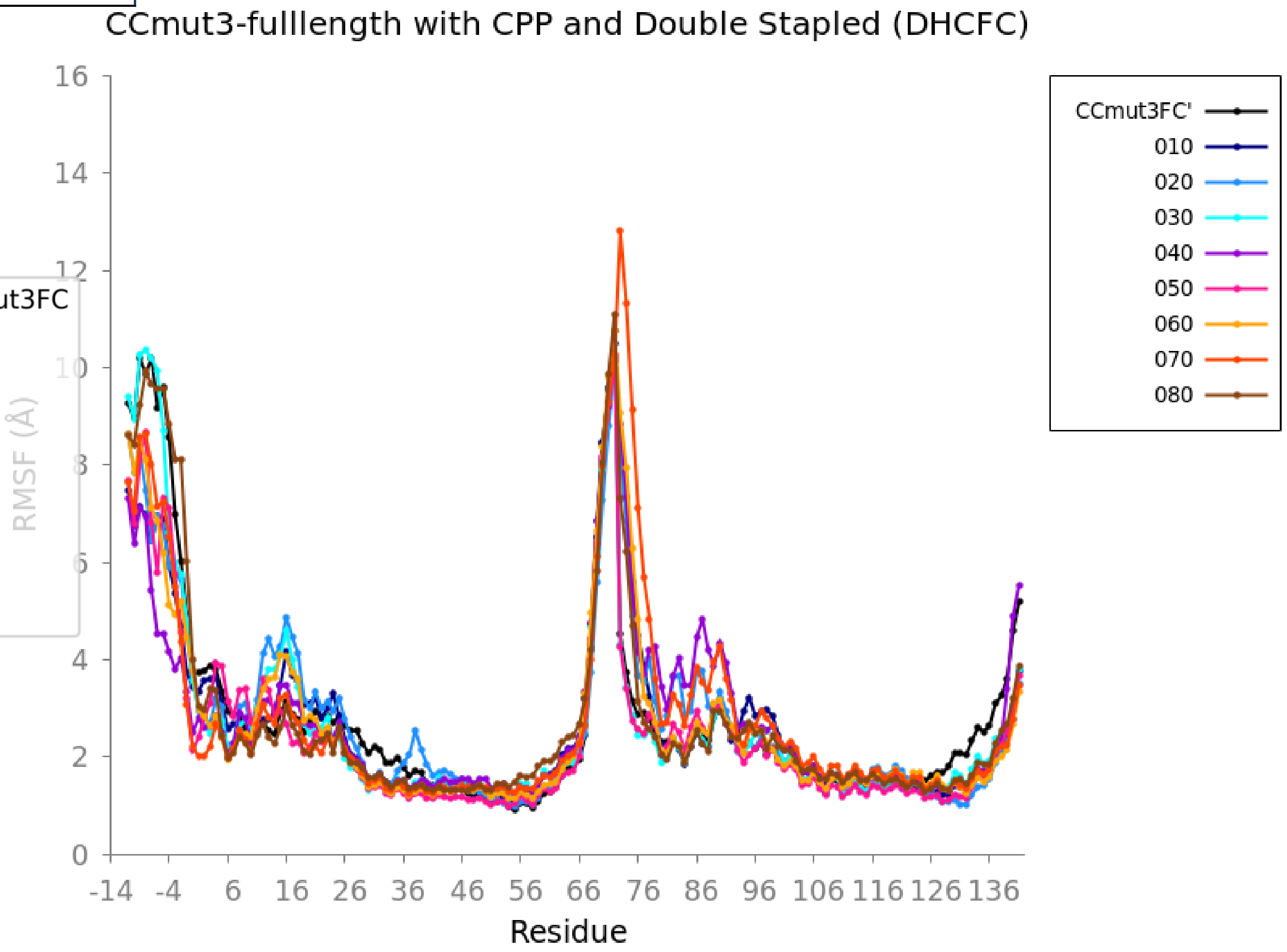

a)

| System | ΔG | Std. Err. of Mean | ΔΔG |
|---|---|---|---|
| DHCT-010 | -37.72 | 0.56 | 6.67 |
| DHCT-020 | -40.54 | 0.64 | 3.85 |
| DHCT-030 | -39.85 | 0.58 | 4.54 |
| DHCT-040 | -26.26 | 0.65 | 18.13 |
| DHCT-050 | -37.80 | 0.54 | 6.59 |
| DHCT-060 | -47.15 | 0.53 | -2.76 |
| DHCT-070 | -43.70 | 0.52 | 0.69 |
| CCmut3T | -44.39 | 0.59 | 0.00 |

b)

| System | ΔG | Std. Err. of Mean | ΔΔG |
|---|---|---|---|
| DHCTC-010 | -56.40 | 0.62 | 2.42 |
| DHCTC-020 | -47.46 | 0.69 | 11.35 |
| DHCTC-030 | -49.88 | 0.62 | 8.94 |
| DHCTC-040 | -45.53 | 0.64 | 13.29 |
| DHCTC-050 | -58.40 | 0.59 | 0.42 |
| DHCTC-060 | -60.02 | 0.65 | -1.21 |
| DHCTC-070 | -57.35 | 0.56 | 1.47 |
| CCmut3TC | -58.82 | 0.65 | 0.00 |

c)

| System | ΔG | Std. Err. of Mean | ΔΔG |
|---|---|---|---|
| DHCF-010 | -57.17 | 0.65 | -4.44 |
| DHCF-020 | -50.51 | 0.72 | 2.23 |
| DHCF-030 | -50.81 | 0.67 | 1.92 |
| DHCF-040 | -48.49 | 0.64 | 4.24 |
| DHCF-050 | -59.89 | 0.59 | -7.16 |
| DHCF-060 | -48.61 | 0.68 | 4.12 |
| DHCF-070 | -53.87 | 0.64 | -1.14 |
| DHCF-080 | -44.27 | 0.66 | 8.46 |
| CCmt3F | -52.73 | 0.67 | 0.00 |

d)

| System | ΔG | Std. Err. of Mean | ΔΔG |
|---|---|---|---|
| DHCFC-010 | -72.18 | 0.63 | 1.70 |
| DHCFC-020 | -59.50 | 0.80 | 14.39 |
| DHCFC-030 | -72.35 | 0.60 | 1.53 |
| DHCFC-040 | -61.83 | 0.70 | 12.05 |
| DHCFC-050 | -79.09 | 0.63 | -5.21 |
| DHCFC-060 | -76.55 | 0.69 | -2.67 |
| DHCFC-070 | -67.99 | 0.67 | 5.89 |
| DHCFC-080 | -75.23 | 0.63 | -1.34 |
| CCmt3FC | -73.89 | 0.50 | 0.00 |

### Salt Bridge – DHCFC

CCmt3FC
DHCFC – 020
DHCFC – 050

LYS NZ – Black vdw
ASP OD1 and OD2 – Orange vdw
GLU OE1 and OE2 – Yellow vdw
ARG NH* - Magenta vdw

| CCmut3FC | | % |
|---|---|---|
| GLU34 | ARG139 | 66.64 |
| GLU46 | ARG127 | 63.92 |
| GLU66 | ARG100 | 39.58 |
| GLU60 | LYS113 | 26.16 |
| GLU32 | LYS141 | 22.11 |

| DHCFC-020 | | % |
|---|---|---|
| GLU34 | ARG129 | 54.31 |
| GLU66 | ARG100 | 37.78 |
| GLU32 | LYS141 | 29.69 |
| GLU46 | ARG127 | 29.09 |
| ASP77 | ARG96 | 23.3 |

| DHCFC-050 | | % |
|---|---|---|
| GLU46 | ARG127 | 79.12 |
| GLU34 | ARG139 | 47.42 |
| GLU60 | LYS113 | 31.18 |
| GLU32 | LYS141 | 29.7 |
| GLU66 | ARG100 | 24.07 |
